# Unveiling the molecular choreography: In-depth single-cell transcriptomic exploration of the regenerative dynamics in stony coral

**DOI:** 10.1101/2024.05.05.592605

**Authors:** Tingyu Han, J.-Y. Chen, Chunpeng He, Zuhong Lu

## Abstract

The coral reef ecosystem faces increasing threats under global climate challenges. One of the core issues is the inability of fragments to quickly grow into a size that can resist environmental pressures in coral transplantation. The observation of accelerated growth during the early stages of coral regeneration provides new insights for addressing this challenge. To investigate related molecular mechanisms, our study focused on the fast-growing stony coral *Acropora muricata* (with chromosome-scale reference genome). Employing diverse techniques, including single-cell RNA sequencing (scRNA-seq), we unveiled related intricate cellular dynamics. Single-cell analysis revealed notable shifts in calicoblasts and epidermal cells around 2-4 weeks post-injury. Gene expression analysis revealed enrichment in immune response and biomineralization pathways. Pseudotime analysis explained the differentiation of epidermal cells into calicoblasts, while time-course analysis identified key genes associated with dynamic biomineralization changes. This study enhances our understanding of coral regeneration, offering insights for protective strategies to foster coral growth.

## Introduction

The coral reef ecosystem is undergoing a transformative phase with increasingly brief intervals between coral bleaching events, impeding the recovery of mature coral colonies. As of the 1880s, Earth’s surface temperature has surged by nearly 1°C due to anthropogenic climate warming and El Niño, continuously setting unprecedented records. Recent global sea surface temperature spikes further accelerate the warming trend, reducing the median return time between severe coral bleaching events to just 6 years by 2016(*1*). This escalating frequency poses a significant threat to coral reefs, vital thermal refuges for diverse marine species(*2*). Ominous predictions suggest these reefs may vanish by midcentury(*1*).

Faced with this challenge, urgent and informed management strategies are crucial to protect coral reefs and restore their compromised structure and ecological functionality. Globally, researchers, conservationists, and environmental managers are actively devising innovative strategies to preserve coral reef ecosystems amidst diverse local and global threats(*3*). Genetic and reproductive interventions, like managed selection and breeding, aim to propagate stress-tolerant corals(*4*). Techniques such as gamete and larval capture(*5*), coral cryopreservation(*6*), and genetic manipulation(*7*) offer novel ways to enhance genetic diversity. Physiological interventions, including pre-exposure(*8*) and algal symbiont manipulation(*9*), tap into corals’ adaptive abilities. Strategies like microbiome manipulation(*10*), antibiotic application(*11*), and phage therapy(*12*)target coral microbial communities for improved health and stress resistance. Coral population interventions, like managed relocation(*13*), propose shifting corals to favorable environments. Environmental strategies, including shading(*14*) and abiotic ocean acidification(*15*) interventions, suggest innovative ways to mitigate thermal stress and modify seawater chemistry. Despite implemented measures, coral reefs decline unabated. A crucial factor is the need for substantial recovery volume, preserving reef structure while mitigating coral losses(*16*). Recognizing this, micro-fragmentation leverages corals’ regenerative capacity, segmenting colonies for transplantation. Research indicates accelerated growth in micro-fragments post-fragmentation(*17, 18*), highlighting its potential efficacy. However, molecular mechanisms remain unclear. Lock et al. (2022)(*19*) proposes that micro-fragmentation in *Porites lobata*disrupts calcium homeostasis, energizing coral expansion as a wound healing response, contributing to survival. The study hypothesizes enhanced calcification rates for skeletal regeneration, but understanding these molecular processes is limited. To uncover the molecular mechanisms driving stony coral regeneration, we focus on *Acropora muricata*, a fast-growing coral crucial for reef structure(*20*). Our study integrates scRNA-seq, bulk RNA-seq, full-length transcriptome sequencing, and miRNA analysis for a holistic view of coral regeneration. Differences between scRNA-seq and bulk RNA-seq data emphasize the complex regulatory processes. Our multifaceted approach aims to elucidate cellular dynamics, gene expression, and communication networks, contributing significantly to coral regeneration understanding. This foundation informs future studies on coral resilience and adaptation, guiding conservation efforts for threatened coral reef ecosystems.

## Results

### Morphological and HRCT 3D reconstruction of coral regeneration

In our exploration of optimal regeneration stages for *A. muricata*, a preliminary experiment revealed a peak regeneration period approximately 2−4 weeks post-injury. Systematically collecting samples from days 0 to 39, with observations every 3 days, morphological studies indicated the most rapid recovery between 15 and 27 days (Fig. 1a). Validating these findings, selected samples from days 0, 14, 21, and 28 underwent high-resolution computed tomography (HRCT) three-dimensional (3D) reconstructions of their truncated areas (Fig. 1b). Results showed that by day 14, the lumen diameter significantly decreased, indicating calcified substance deposition. By day 21, the deposition increased, completely filling the lumen, suggesting enhanced biomineralization. By day 28, the lumen returned to a morphology closer to day 14, signifying the late repair stage. The regenerative capacity of coral is notably robust around day 21, prompting the selection of samples from this time point for subsequent single-cell level bioinformatics analysis, comparing them with normally growing coral samples.

**Fig. 1.**
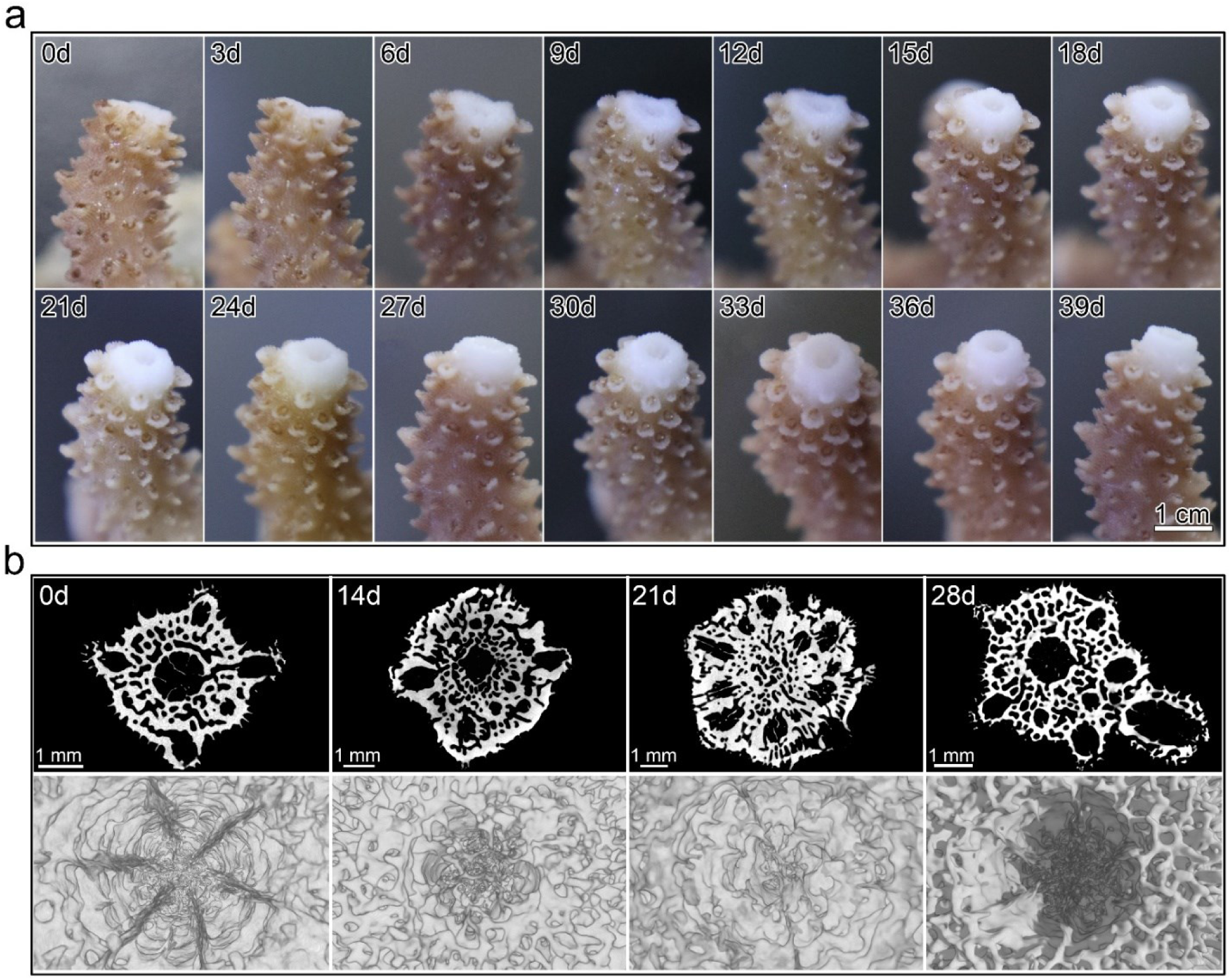
Monitoring coral regeneration growth and HRCT 3D reconstruction. **a**, Growth status every 3 days after coral damage. **b**, 3D reconstruction of cross-sectional views during four stages of coral damage repair using HRCT.

### Establishment of the *A. muricata* cell atlas utilizing 10x Genomics

We analyzed 9,053 filtered cells from scRNA-seq datasets of healthy and regenerating *A. muricata* coral. Using established cell markers, we identified 13 cell types, including gastrodermal cells, neurons, calicoblasts, alga-hosting cells, epidermal cells, cnidocytes, gland cells, progenitor cells, immune cells, muscle cells, digestive filament cells, and two unassigned cell clusters (Supplementary Fig. 1 and Supplementary Dataset 1). Subsequent re-clustering of the 11 function-specific cell clusters was conducted for further analysis (Fig. 2a, b; Supplementary Fig. 2−8; Supplementary Dataset 2).

**Fig. 2.**
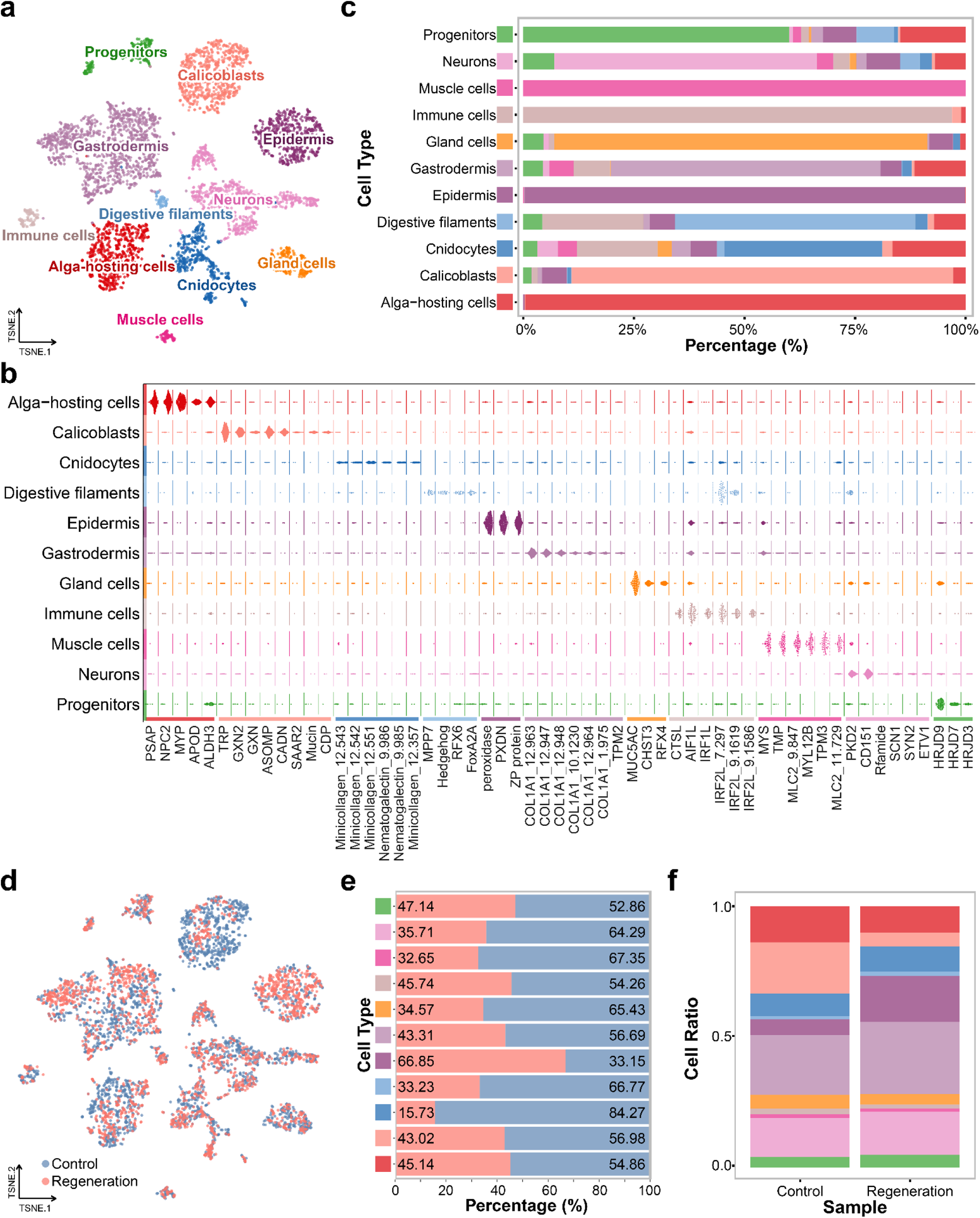
Single-cell atlas of coral regeneration. **a**, t-SNE visualization of integrated analysis for two coral samples. **b**, Beeswarm plots displaying log-transformed UMI counts of marker genes in each cell type. **c**, Reliability assessment of cell clustering. **d**, t-SNE visualization of different coral samples. **e**, Comparison of the proportion of identical cell types in different samples. **f**, Distribution of cell types in different samples.

To validate cell clustering reliability, we used UCell to score Seurat objects based on cell markers and visualized cell type composition within clusters (Fig. 2c). Most clusters were predominantly composed of cells from their respective types, except for cnidocytes and digestive filaments. Immune cells were notably abundant in both cnidocytes and digestive filament cells, suggesting potential immune-related gene or process involvement. Considering cnidocytes’ role in prey capture and anti-predator defense(*21*), and digestive filaments’ function in food acquisition(*22*), these cell types may exhibit immune functions.

To compare cell distribution between samples, we generated a t-SNE plot (Fig. 2d), with “Control” representing the healthy sample and “Regeneration” representing the vigorously regenerating sample. Both samples exhibited the 11 identified cell types. Despite total cell number variations, the relative proportions and distribution of different cell types within and across both samples remained consistent (Fig. 2e and f). Calicoblasts and epidermal cells showed notable differences in cell numbers between samples, suggesting their specific involvement in the regeneration process. These findings indicate that specific cell types may adjust during coral regeneration to meet functional and physiological demands.

To identify additional cell type-specific markers, we analyzed highly expressed genes conserved across conditions. Notably, certain genes, distinct from those previously mentioned, exhibited significant upregulation within the same cell types across both samples (Supplementary Fig. 9 and Supplementary Dataset 3). Sulfate transporter(*23*) and saccharopine dehydrogenase(*24*) showed specific high expression in alga-hosting cells, supporting earlier reports of elevated levels in symbiotic hosts. Unexpectedly, *Ca2*, known for its role in biomineralization, showed heightened expression in alga-hosting cells. This could be due to its additional crucial role in carbon-concentrating mechanisms essential for symbiont photosynthesis(*25*). Leucine-rich repeat extensin-like proteins showed specific high expression in cnidocytes, consistent with Levy et al. (2021)’s findings(*26*). Immune cells displayed specific high expression of RIG-I-like receptor, an antiviral protein linked to the innate immune response(*27*). Furthermore, a set of genes associated with calcium ion binding (GO:0005509), encompassing calretinin, neurocalcin homolog, and calmodulin, were specifically highly expressed within neurons. These findings unveil a range of potential cell markers that contribute to our understanding of the functional roles and specialized characteristics of diverse cell types within coral organisms.

### Exploring differential gene expression patterns across conditions

Utilizing the Wilcoxon rank-sum test, we compared single-cell transcriptomic data between healthy and regenerating coral samples, identifying 536 significantly differentially expressed genes (DEGs) (Supplementary Dataset 4). These DEGs were distributed across various cell types, with notable variations in the number of upregulated and downregulated genes within each category (Fig. 3a). Immune and muscle cells did not show significant DEGs in this analysis. 9 DEGs upregulated in at least two cell types, 69 DEGs upregulated in a single cell type. Conversely, 79 DEGs downregulated in at least two cell types, while 233 DEGs downregulated in a single cell type.

**Fig. 3.**
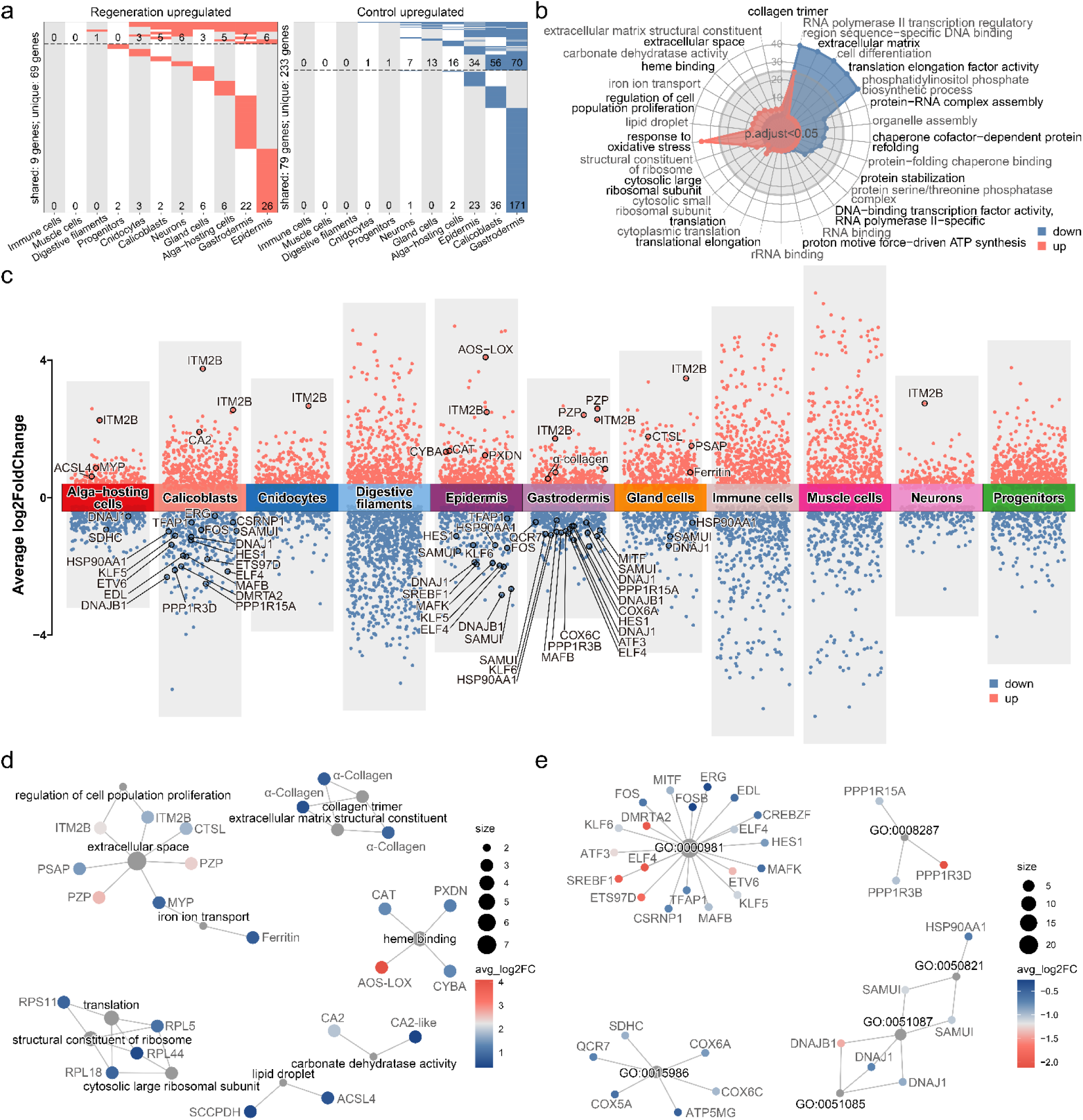
Analysis of differentially expressed genes in coral regeneration. **a**, Heatmaps illustrating the distribution of significantly DEGs, with numbers below the dotted lines indicating specific DEGs in only one cell type, and numbers above the dotted lines indicating DEGs in at least two cell types. **b**, Radar plot depicting GO enrichment pathways of significantly DEGs. **c**, Manhattan plot for significantly DEGs. **d**, Cnetplot displaying the network linking significantly upregulated DEGs to enriched GO pathways. **e**, Cnetplot displays a partial network linking significantly downregulated DEGs to enriched GO pathways, with the remaining part shown in Supplementary Fig. 10.

Further analysis of upregulated and downregulated DEGs revealed significant enrichment in 13 and 21 Gene Ontology (GO) pathways, respectively (Fig. 3b and Supplementary Dataset 5). Among the upregulated DEGs, the GO pathway associated with the extracellular space (GO:0005615) exhibited the highest number of enriched genes. Notably, *Itm2b*, which was also associated with the regulation of cell population proliferation (GO:0042127), exhibited remarkable upregulation in seven distinct cell clusters (Fig. 3c and d). Previous research has highlighted genes containing a conserved BRICHOS domain (PF04089), such as *Itm2b*, as being differentially expressed in corals under heat and acidification stress conditions, suggesting a potential role in the coral response to environmental stress(*28*). Immune-related genes like *Ctsl*, *Psap* (which may play a role in preventing oxidative stress(*29*)), and pregnancy zone proteins (PZPs, identified in the skeletal proteome of the sea star *Patiria miniate*(*30*)) were also enriched in this pathway (Fig. 3c and d). The epidermal cells showed enrichment in the heme binding pathway (GO:0020037), including genes like *allene oxide synthase-lipoxygenase* (*AOS-LOX*), *Cat*, *Cyba*, and *peroxidasin homolog* (*Pxdn*) (Fig. 3c and d). These genes are classified as oxidoreductases. The AOS-LOX pathway is capable of producing oxylipins, which are crucial stress mediators that indirectly regulate the expression of defense genes. Lohelaid et al. (2014) has demonstrated an upregulation of the AOS-LOX pathway in response to coral wounding, indicating its role in coral damage response and regeneration, akin to its function in plants(*31*). CAT is an enzyme that catalyzes the degradation of hydrogen peroxide, scavenging reactive oxygen species (ROS), and promoting cell growth, playing a crucial role in the coral response to external stimuli(*32*). CYBA is one of the subunits of NADPH oxidase (NOX), involved in ROS production and playing an essential role in the immune system, and it has been observed upregulated in response to coral fragmentation(*19*). PXDN is another important immune response gene, with reports of significant upregulation in response to stony coral tissue loss disease(*33*) and heat stress(*34*).

DEGs related to collagen trimer (GO:0005581) and extracellular matrix (ECM) structural constituent (GO:0005201) were predominantly concentrated in the gastrodermis (Fig. 3c and d). These genes belong to the alpha collagen group, a key component of the extracellular matrix critical for regulating wound healing phases(*35*). In calicoblasts and alga-hosting cells, there was significant upregulation of *Ca2* associated with carbonate dehydratase activity (GO:0004089). Notably, this upregulation was more pronounced in calicoblasts, suggesting a potential role in promoting coral regeneration (Fig. 3c and d). Genes involved in iron ion transport (GO:0006826) included ferritin in gland cells, which plays a crucial role in maintaining iron homeostasis and protecting against oxidative damage caused by free iron and ROS(*36*). And alga-hosting cells exhibited major yolk protein (MYP), a transferrin-like protein derived from digestive filaments. This protein is known to serve as a source of nutrients and energy(*37*) or participate in the transport of zinc derived from food(*38*) (Fig. 3c and d). DEGs enriched in lipid droplet (GO:0005811) were identified in alga-hosting cells, including *long-chain-fatty-acid--CoA ligase 4* (*Acsl4*), involved in fatty acid metabolism and providing critical components for energy storage and nutrient availability in corals(*39*). Additionally, *saccharopine dehydrogenase-like oxidoreductase* (*Sccpdh*) was found, serving as a candidate marker of coral alga-hosting cells (Fig. 3c and d).

For the downregulated DEGs, a group primarily located in the gastrodermis exhibited enrichment in multiple GO terms, particularly in ribosomal proteins and elongation factors, essential components of ribosomes and cyclically associated with ribosomes during protein synthesis elongation (Supplementary Fig. 10). The observed decrease in their expression in the gastrodermis suggests a potential redirection of cellular resources towards other vital processes associated with coral regeneration, such as tissue repair, energy allocation, or responses to stress. Another group of DEGs related to DNA-binding transcription factor activity, RNA polymerase II-specific (GO:0000981), was prominently expressed in the epidermal cells, calicoblasts, and gastrodermal cells (Fig. 3c and e). DEGs involved in proton motive force-driven ATP synthesis (GO:0015986), primarily composed of components of the cytochrome c oxidase complex, displayed downregulation in the gastrodermis (Fig. 3c and e). Cytochrome c oxidase is a crucial enzyme complex responsible for the final step of the electron transport chain in mitochondria, a process essential for generating cellular energy in the form of ATP(*40, 41*). The observed downregulation of these DEGs may suggest a metabolic shift, possibly reflecting altered energy demands or changes in mitochondrial function in the gastrodermis in response to the coral regenerative process. Additionally, a group of DEGs related to protein folding, including *heat shock protein HSP 90-alpha* (*Hsp90aa1*) and *DnaJ protein homolog 1* (*Dnajb1*), showed significant downregulation in the gastrodermal cells, calicoblasts, epidermal cells, and gland cells (Fig. 3c and e). Finally, a set of protein phosphatase 1 regulatory (PPP1R) subunits derived from the gastrodermal cells and calicoblasts were enriched in GO:0008287, which relates to the protein serine/threonine phosphatase complex (Fig. 3c and e).

### Changes in cell–cell communication networks during coral regeneration

To gain deeper insights into the intricate cell–cell communication networks orchestrating the repair process, we conducted a comparative analysis using CellChat. The results unveiled significant associations of intercellular communication in both states of coral cells with the ECM–receptor regulatory network, implicating key ECM proteins such as collagen, thrombospondin (THBS), heparan sulfate proteoglycan (HSPG), and laminin, play pivotal roles in modulating cell–cell and cell–matrix interactions(*42–45*) (Supplementary Dataset 6.1−3). Noteworthy differences in interactions among these pathways emerged during coral regeneration, with increased interactions directed towards digestive filaments and muscle cells, and decreased interactions targeting calicoblasts, epidermal cells, and gastrodermal cells (Fig. 4a). The interaction strength between gastrodermal cells and calicoblasts witnessed a pronounced decline. Mapping the inferred intercellular communication networks onto a shared two-dimensional manifold revealed clustering based on functional similarity, emphasizing the indispensability of identical signaling pathways for both normal coral growth and the repair process, therefore they might not be pivotal factors promoting regeneration during the repair process (Fig. 4b). Additionally, the overall communication probability between the two conditions showed that THBS and laminin were highly active in normal samples (Fig. 4c).

**Fig. 4.**
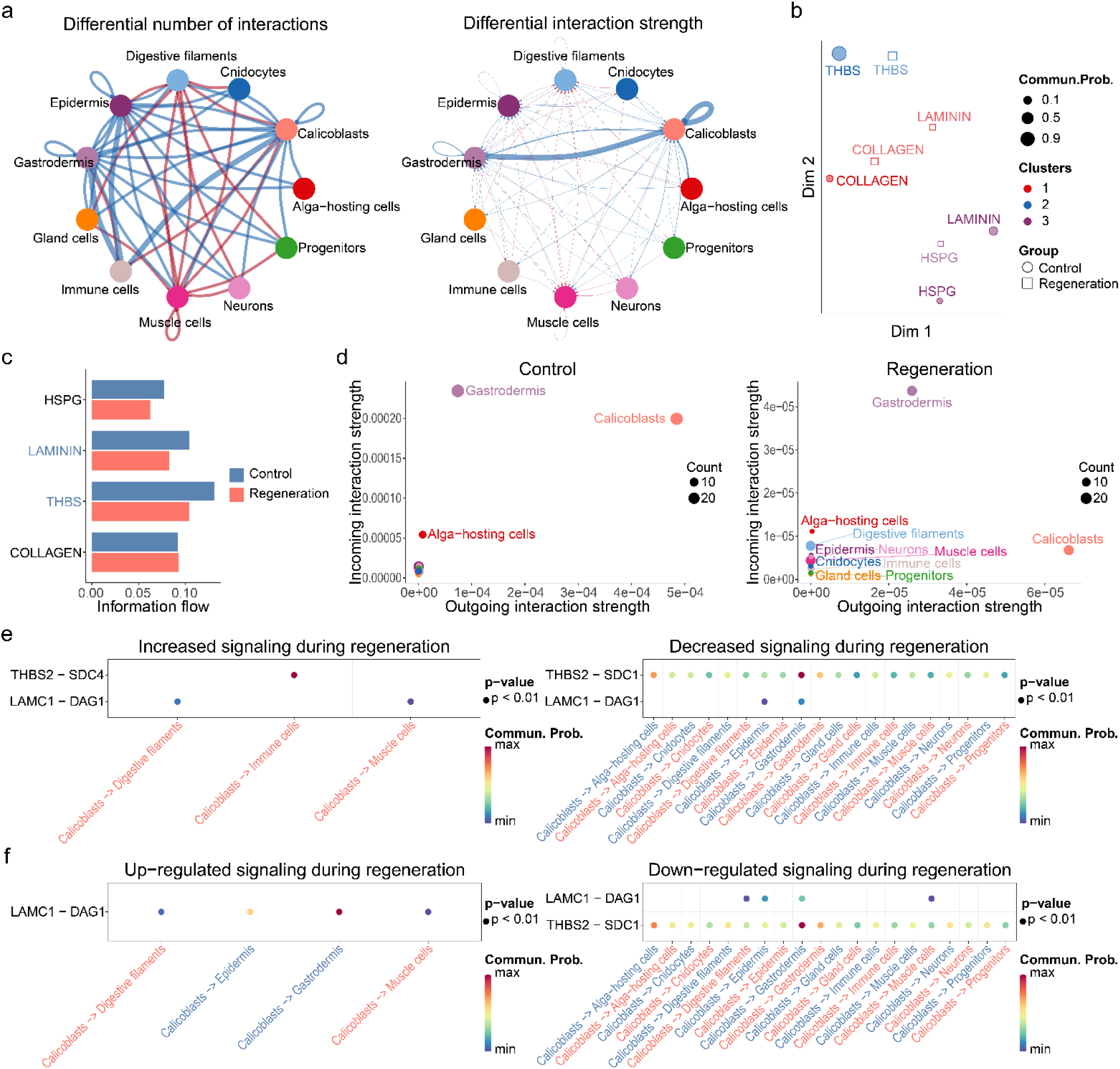
Cell–cell communication networks during coral regeneration. **a**, Visualizing the differential number of interactions or interaction strength in the cell-cell communication network between two datasets. Edges colored in red (or blue) indicate enhanced (or reduced) signaling in the regeneration dataset compared to the control dataset. **b**, Two-dimensional diagram of the signaling network clustered based on functional similarity. **c**, Comparison of overall information flow in each signaling pathway across different samples, with pathways highlighted in blue text enriched in the control sample. **d**, Visualization of two-dimensional diagrams depicting the outgoing and incoming interaction strengths across different cell types. **e**, Increased and decreased in the communication probability of signaling ligand-receptor pairs from calicoblasts to other cell types in the regeneration sample compared to the control sample. **f**, Up-and down-regulated in the communication probability of signaling ligand-receptor pairs from calicoblasts to other cell types in the regeneration sample compared to the control sample.

In-depth analysis of outgoing and incoming interaction strength pinpointed calicoblasts as the cell population exhibiting the most significant changes in sending or receiving signals during coral regeneration (Fig. 4d). Building upon this, we investigated the signaling ligand-receptor pairs directed from calicoblasts towards other cell types (Fig. 4e, f, and Supplementary Dataset 6.4). The results revealed heightened and upregulated LAMC1-DAG1 signaling from calicoblasts to digestive filaments and muscle cells (Supplementary Fig. 11a). Additionally, a surge in THBS2-SDC4 signaling was observed towards immune cells, while THBS2-SDC1 signaling towards multiple cell types diminished (Supplementary Fig. 11b). Conversely, signaling ligand-receptor pairs targeting calicoblasts from other cell types showed no enhancement in signals but identified weakened and downregulated COL1A2-SDC1 signaling from gastrodermal cells to calicoblasts (Supplementary Fig. 12, 13, and Supplementary Dataset 6.4). These findings highlight the intricate recalibrations in cell-cell communication networks during coral regeneration, emphasizing the pivotal role of specific signaling pathways and the differential responses of various cell types.

### Differentiation of epidermis during coral regeneration

In stony coral development, certain epidermal cells transform into calicoblasts, initiating rapid skeletal deposition(*26*). Pseudotime analysis was employed to explore the relationship and potential transitions between these cell types during coral regeneration (Fig. 5). Results showed that, during normal coral development, some epidermal cells transitioned into two distinct calicoblast types (Fig. 5a−c). In contrast, during regeneration, more epidermal cells predominantly transformed into a single subtype of calicoblasts (Fig. 5d), suggesting that the primary calicoblast types involved in biomineralization differ between normal coral growth and the repair process, highlighting the dynamic nature of coral cell differentiation and function in response to different developmental and regenerative cues. The observed subtype may also represent a transitional state, potentially reverting to normal development and transforming into another subtype in later regeneration stages. We constructed a heatmap and performed GO enrichment analysis to explore the dynamic expression changes of genes associated with cell transformation (Fig. 5e and Supplementary Dataset 7). Under normal conditions, epidermal cells exhibit three distinct peaks during development, suggesting different developmental stages. This aligns with pseudotime analysis results, although epidermal cells at different developmental stages are clustered together (Supplementary Fig. 14). Calicoblasts under normal conditions show a relatively uniform distribution of cell proportions from the initial state to maturity. During regeneration, both epidermal cells and calicoblasts in earlier developmental stages show a significant increase in cell numbers, suggesting an enhanced capacity to differentiate more epidermal cells into calicoblasts during coral repair. This implies that the biomineralization potential of calicoblasts in the early developmental stages may differ from that of mature stages.

**Fig. 5.**
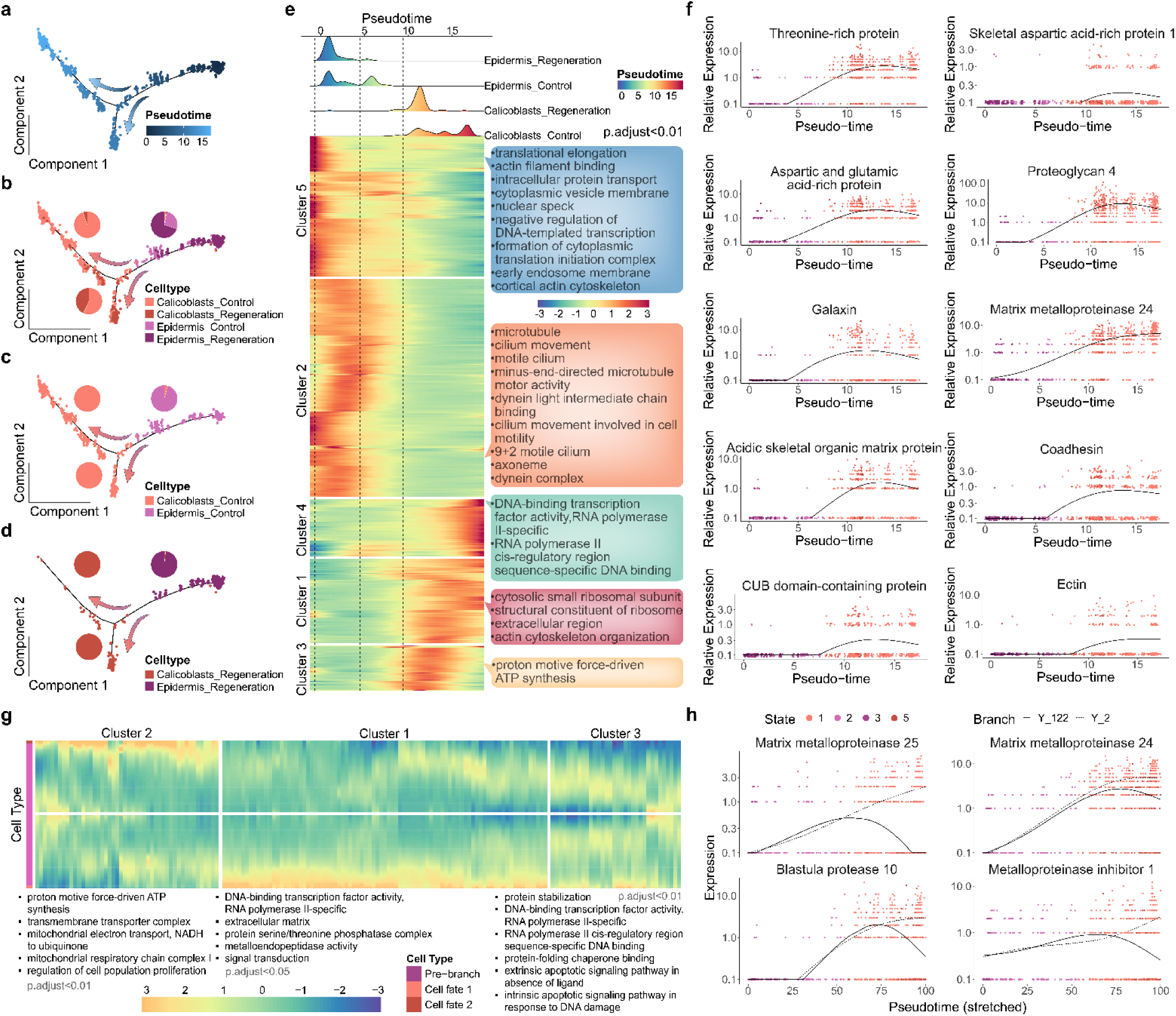
Pseudotime trajectory analysis during coral regeneration. **a**, Pseudotime analysis of two cell types across both samples, with colors ranging from dark to light indicating developmental progression from early to late stages. **b-d**, Cellular distribution of the two cell types along the cell trajectory in all samples, control sample, and regeneration sample, respectively. **e**, Ridge plot and heatmap visualization of the pseudotime analysis. The upper ridge plot illustrates the pseudotime distribution of cell numbers for the two cell types across different samples. The lower heatmap displays clustered analysis of gene expression related to intercellular state transitions and corresponding GO functional enrichment analysis. **f**, Pseudotime analysis of biomineralization-related genes based on cell types. **g**, Heatmap displaying different fates and clustered analysis of DEGs associated with epidermal cell to calicoblast differentiation, along with corresponding GO functional enrichment analysis. **h**, Differential expression analysis of genes enriched in GO terms (extracellular matrix and metallopeptidase activity) in Cluster 1 from panel g.

During early calicoblast development, specifically in cluster 1 and cluster 3, several genes exhibited distinct upregulation, prominently enriched within the extracellular region (GO:0005576), with proteins like threonine-rich protein(*46*), aspartic and glutamic acid-rich protein, galaxin, acidic skeletal organic matrix protein, CUB domain-containing protein (*Cdcp1*), and skeletal aspartic acid-rich protein 1 (*Saarp1*)(*47, 48*) (Fig. 5f and Supplementary Dataset 7). These proteins are known components of the coral skeleton. Moreover, we also found other genes potentially involved in coral biomineralization that were specifically upregulated in cluster 1. Notably, proteoglycan 4 (PRG4) is a key protein, as existing studies suggest that the biomineralization process begins with the secretion of a proteoglycan matrix(*49*), further confirming that the above-mentioned genes in cluster 1 are specifically highly expressed in early-stage calicoblasts. Matrix metalloproteinase 24 (MMP24), belonging to the matrix metalloproteinase gene family which plays a vital role in balancing and regulating the degradation of ECM proteins(*50*), might also participate in coral calcification. Additionally, we identified AOS-LOX, peroxinectin, and melanotransferrin, which associated with immune response(*51, 52*), showing similar expression patterns (Supplementary Fig. 15a and Supplementary Dataset 7). Cluster 3 showed specific upregulation of three genes (*galaxin 2*, *Saarp2*, and *Mucin*) linked to biomineralization (Supplementary Fig. 15b and Supplementary Dataset 7). In the advanced developmental phases of calicoblasts within cluster 4, limited genes, including *uncharacterized skeletal organic matrix protein 4* (*Usomp4*), *Mmp25*, and *metalloproteinase inhibitor 1* (*Timp1*), exhibited distinct high expression associated with biomineralization (Supplementary Fig. 15c and Supplementary Dataset 7). These findings underscore the potential variability of key genes responsible for biomineralization in calicoblasts at different developmental stages, and this variance might be intricately linked to the underlying mechanisms governing coral skeletal formation.

During the transformation of coral epidermal cells into calicoblasts, two distinct fates were observed. To delineate the disparities between these fates, we conducted a GO enrichment analysis (Fig. 5g and Supplementary Dataset 8). Genes associated with cell fate 1 (state 1), primarily composed of calicoblasts in a normal state, were significantly upregulated in two clusters (1 and 3). Cluster 1 exhibited enrichment in extracellular matrix and metalloendopeptidase activity, including genes such as *Mmp24*, *Mmp25*, *blastula protease 10* (*Bp10*), and *Timp1* (Fig. 5h). Cluster 3 showed enrichment in processes related to protein stabilization and apoptotic signaling pathways, involving genes like *Hsp90aa1*, *Dnajb1*, *Bcl2*, and *apoptosis regulator R1* (*Ar1*) (Supplementary Fig. 16). Genes associated with cell fate 2 (state 5), predominantly composed of calicoblasts in a regenerating state, were enriched in proton motive force-driven ATP synthesis, transmembrane transporter complex, and regulation of cell population proliferation (Supplementary Fig. 17). Additionally, we identified a set of significantly upregulated genes associated with biomineralization in cell fate 2, including *Ca2*, *Saarp2*, *galaxin*, *galaxin2*, and *Mucin*. In cell fate 1, we observed the upregulation of *Usomp4*, *Cdcp2*, and *calumenin* (Supplementary Fig. 18). These results highlight differential gene expression during epidermal cell transition to calicoblasts and underscore distinctions in gene expression between calicoblasts in normal and regenerative states. This indicates substantial changes in gene expression throughout development, suggesting mature calicoblasts may exhibit unique patterns and functions compared to early-stage calicoblasts. Furthermore, this supports the earlier hypothesis that the two calicoblast clusters playing different roles in coral biomineralization may result from temporal differences in developmental stages rather than simultaneous differentiation into cell types with diverse functionalities.

### Time-course analysis of single-cell level DEGs in bulk RNA-seq data

The preceding contents have provided a thorough exploration of the differences in single-cell transcriptomics between healthy and regenerating coral samples. However, an intriguing question remains unanswered – whether these observed differences extend to the broader transcriptomic landscape and exhibit consistent patterns across various time points during the repair process. To address this knowledge gap, we broaden our focus to the transcriptome-wide scale, aiming to investigate whether the differences observed at the single-cell level are reflected in the bulk RNA-seq data. Upon a comprehensive examination, it appears that the dynamic expression changes of DEGs and biomineralization-related genes identified in the single-cell transcriptome do not exhibit significant patterns in bulk RNA-seq data (Fig. 6 and Supplementary Dataset 9). To gain deeper insights into the sustained effects of these genes during coral repair, we employed the TCseq method for clustering based on temporal gene expression patterns. The results indicate that genes significantly up-or down-regulated in the single-cell transcriptomes do not uniformly exhibit similar changes in expression in the bulk RNA-seq data at the same time point (Fig. 6a and b).

**Fig. 6.**
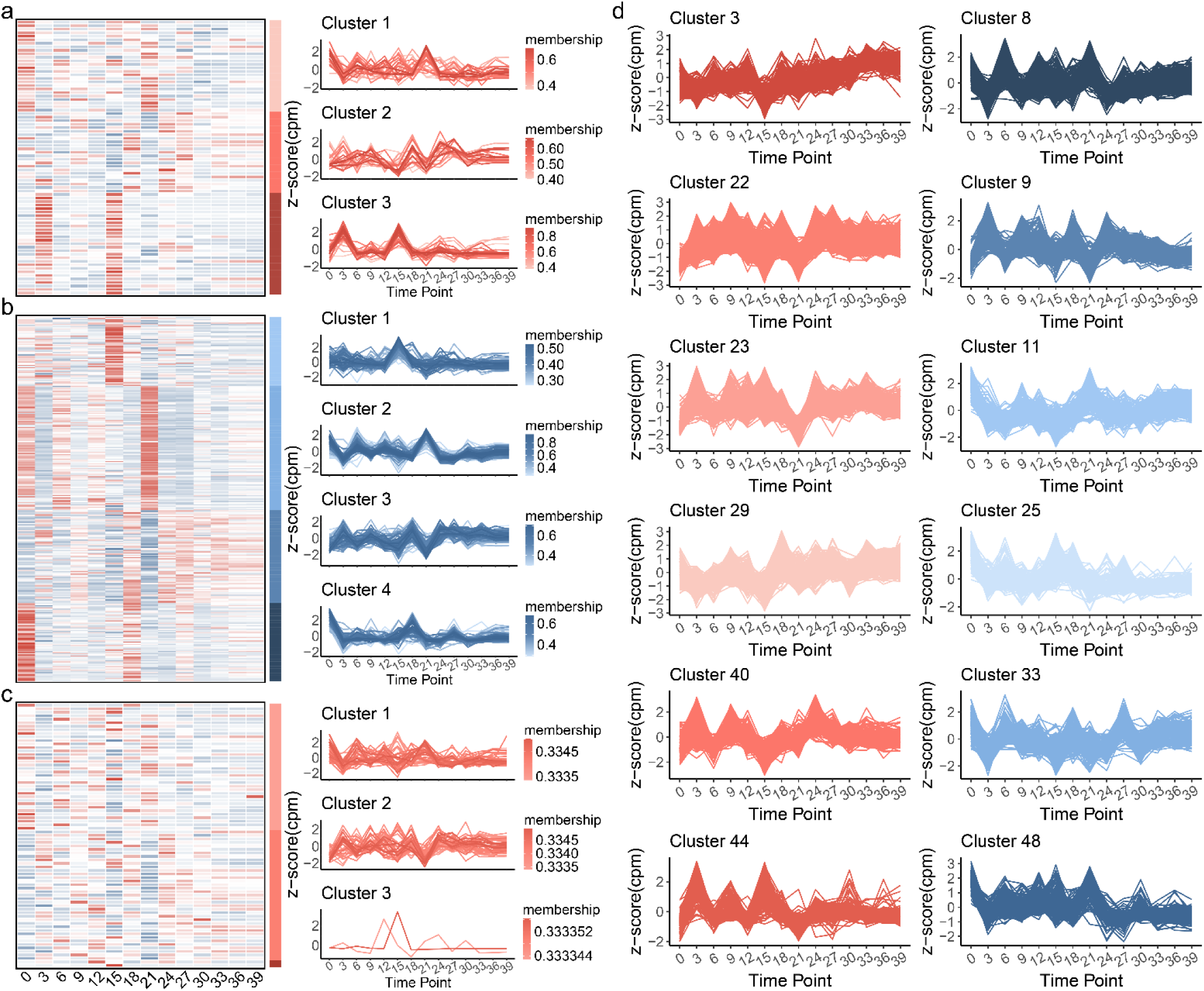
Temporal analysis of DEGs in bulk RNA-seq data. **a-c**, Temporal analysis of significantly upregulated, downregulated, and biomineralization-related DEGs at the single-cell level, respectively. On the right are the temporal expression clustering results obtained using the TCseq method, and on the left are corresponding heatmaps for different clusters. **d**, Significant temporal expression patterns of all bulk RNA-seq DEGs analyzed using the STEM method.

For DEGs showing significant upregulation in the single-cell transcriptome, we categorized them into three clusters based on their temporal gene expression patterns (Fig. 6a and Supplementary Dataset 9.2). DEGs in cluster 1 exhibit significant downregulation in the early stages of coral repair, followed by irregular fluctuations over time, with the majority peaking at day 21. Key members include genes possibly associated with biomineralization, such as *Pzp*, *calmodulin*, *Ca2*, and *calumenin*, along with stress-related genes like *Pxdn*. DEGs in cluster 2 display continuous fluctuation during coral repair with an overall increasing trend until reaching relative stability in the later stages, including a group of alpha collagens. DEGs in cluster 3 show significant peaks in expression at the early (day 3) and mid (day 15) stages, encompassing stress-related genes (*Itm2b*, *AOS-LOX*, *Ctsl*, and *ferritin*) and symbiosis-related genes (*Sccpdh* and *Myp*).

For DEGs exhibiting downregulation in the single-cell transcriptome, we clustered them into four groups (Fig. 6b and Supplementary Dataset 9.3). DEGs in cluster 1 display a noticeable peak at day 15, including stress-related genes (*soma ferritin* and *Hsp90aa1*) and biomineralization-related genes (*Mmp25*, *Timp1*, *Prg4*, and *Bp10*). DEGs in cluster 2 exhibit a similar expression pattern to the first type of temporal gene expression pattern observed in the upregulated genes in the single-cell transcriptome, primarily involving a set of ribosomal proteins, ATP synthase-related genes, cytochrome c oxidase, and a biomineralization-related gene (*Usomp4*). Interestingly, this contradicts the results at the single-cell level, where these genes are significantly downregulated in the regenerating coral’s single-cell transcriptome at day 21 but significantly upregulated in the corresponding bulk RNA-seq data. DEGs in cluster 3 show significant downregulation at days 15 and 21, while DEGs in cluster 4 exhibit significant downregulation in the early stages of coral repair, followed by varying degrees of recovery at days 18 and 27, including *Mmp24* and *Timp4*.

Concerning biomineralization-related genes, we categorized them into three classes (Fig. 6c and Supplementary Dataset 9.4). The results reveal that genes involved in calcification vary across different stages of coral repair. However, their significant fluctuations are more pronounced before the mid-stage of regeneration, subsequently stabilizing in the later stages. This heightened activity during the initial phases may suggest a crucial role in the early processes of biomineralization and skeletal formation. As the repair progresses to later stages, the expression levels of these genes become more stabilized, indicating a potential shift in focus or a reduced requirement for their active participation in the later phases of coral regeneration.

To further explore temporal gene expression patterns during coral repair, STEM software identified 12 clusters with significant temporal features based on genes showing significant differential expression at least at one time point (Fig. 6d, Supplementary Fig. 19, and Supplementary Dataset 9.5). Gene expression changes showed a noticeable reduction in magnitude in the later stages of the study. Among the USOMPs, clusters 3, 11, 22, 40, and 48 exhibited distinct expression patterns. Notably, *Usomp4* (in cluster 11) and one isoform of *Usomp5* (in cluster 48) displayed significant downregulation in the early repair stages, with a partial recovery but consistently lower expression levels than the normal state throughout the study period. Similarly, *Usomp7* and *Usomp1* (both in cluster 11) showed significant downregulation on the third day post-injury, maintaining consistently lower expression levels thereafter. In contrast, only *Usomp3* (in cluster 3) and another isoform of *Usomp5* (in cluster 22) displayed upregulation during certain periods of coral repair. *Galaxin 2* (in cluster 29) remained significantly upregulated in the middle to later stages. The expression of acidic skeletal organic matrix protein (in cluster 3) and *Saarp1* (in cluster 22) fluctuated in the early stages but maintained relatively high expression levels in the later stages. Particularly, *Ca2* (in cluster 2) consistently and significantly increased in fluctuations, with high expression levels throughout the study.

In addition, we utilized PacBio sequencing technology to conduct comprehensive full-length transcriptome sequencing on coral samples collected on the 21st day of regeneration and from normal coral specimens. Intriguingly, our analysis did not reveal substantial distinctions in critical aspects such as alternative splicing, alternative polyadenylation, and fusion genes between these specified samples. In tandem, a meticulous examination through miRNA sequencing failed to discern any noteworthy differences in this particular domain.

In summary, these results, in conjunction with the preceding content, provide a comprehensive exploration of the transcriptomic landscape in coral samples undergoing regeneration. The investigation spans from single-cell transcriptomics to bulk RNA-seq data, encompassing various time points during the repair process, shedding light on the intricate dynamics of the coral transcriptome during the repair process, offering valuable information for understanding the molecular mechanisms underlying coral regeneration.

## Discussion

The expeditious regeneration of coral colonies is essential for their survival, as demographic indicators, including growth, fecundity, and mortality, exhibit a pronounced reliance on size(*53*). This implies a prioritization of energy allocation towards recovery (regeneration) rather than reproduction in long-lived organisms like corals. Moreover, coral colonies demonstrate a higher degree of integration than conventionally perceived(*54*). Acute mechanical damage, such as tropical storms, fish collisions and predation, and transplant fragmentation can lead to increased mortality rates, and even loss of reproductive capacity(*55*). Previous research has shown a widespread decline in reproductive output in locations distal to the damaged area(*54*), suggesting that injury repair in corals may necessitate extended colony integration, challenging the notion that the energy requirements for this crucial process are solely provided by polyps directly adjacent to the damaged area. Therefore, facilitating the rapid and substantial growth of transplanted corals becomes imperative. This not only reduces baseline costs by minimizing transplant inputs but also ensures the swift attainment of a sufficiently large size to guarantee the structural soundness of the coral colony, providing defense against pathogens and protection from competitors.

Here, our morphological study identified a peak phase in the regeneration of *A. muricata*, occurring approximately 2−4 weeks post-injury. Building upon this observation, we established the *A. muricata* cell atlas using scRNA-seq to comprehensively characterize the cellular dynamics and molecular processes underlying the regeneration phenomenon. The findings demonstrated a consistent repertoire of cell types between normal and regenerating samples, with no identification of novel cell subtypes. This alignment is rational, given that injury repair diverges from physiologic lesions, encompassing tissue regeneration, cell division, proliferation, and the biomineralization process. Nevertheless, discernible adjustments in the quantity of specific cell types during regeneration were noted, particularly an increase in calicoblasts and a decrease in epidermal cells. Developmental trajectory analysis disclosed a continual transformation of epidermal cells into calicoblasts during coral regeneration. In the early stages of calicoblast development, a noteworthy upregulation of the *Ca2* gene was observed, responsible for supplying essential carbonate ions for calcification(*25*). Additionally, a significant augmentation in the secretion of skeletal organic matrix proteins, pivotal components of the coral skeleton(*56*), and metalloproteinases regulating extracellular matrix components(*50*) was evident. Temporal gene expression analysis suggested an eventual stabilization of the overall gene expression pattern in corals after a defined repair period, indicating a return to normal growth conditions. Building upon this, we hypothesize that the accelerated growth during short-term coral regeneration stems from the generation of a substantial quantity of newly formed calicoblasts endowed with high biomineralization capability. This provides an explanation for the immediate short-term accelerated growth observed in coral micro-fragments following initial fragmentation and tissue damage(*17, 18*). However, the precise mechanisms governing the transformation of epidermal cells into calicoblasts remain elusive. Regarding the question of why corals accelerate the deposition of their skeletons in the early stages of repair, similar to other organisms, the coral skeleton plays a crucial role as an attachment substrate for the soft and vulnerable polyp. It acts as the primary physical barrier to protect the soft tissue from external environmental influences (*57*). Simultaneously, due to its richness in peptide neurotoxins, the coral skeleton provides biochemical protection(*58*). Therefore, the observed brief period of rapid growth post-injury serves as a defensive strategy, reinforcing damaged tissues.

In addition, epidermal cells exhibited specific upregulation of a group of oxidoreductases during coral regeneration, with the most prominent being AOS-LOX, a pathway present across all cnidarian lineages(*59*). Previous studies have indicated a significant upregulation of AOS-LOX in tissues near the wound and distal parts of coral colonies in the soft coral *Capnella imbricata* after injury, reaching peak expression at 1 h and 6 h post-injury, respectively(*31*). This suggests that the AOS-LOX pathway serves as a rapidly initiated stress response to acute mechanical damage. However, in our study, by integrating the temporal gene expression patterns of AOS-LOX (Supplementary Dataset 9), we observed significant up-and down-regulation at multiple time points during coral repair, particularly showing early-stage upregulation until later stages where expression stabilized. This implies that AOS-LOX may function as an enzyme with prolonged activity throughout the coral repair process. Because the activity of the enzyme can be influenced by various factors, even if gene expression is downregulated, previously synthesized enzyme molecules may persist and maintain activity for a certain period. Therefore, the intricate temporal regulation of AOS-LOX suggests its potential as a long-acting enzyme during coral regeneration. Several other oxidoreductases, including CAT, CYBA, and PXDN, also exhibited distinct regulation at certain time points during coral injury. Therefore, forthcoming studies should incorporate assessments of enzyme activity to holistically comprehend the functional roles of these enzymes in response to coral wounding.

In our investigation of the rapid regeneration of *A. muricata*, we have uncovered critical insights while acknowledging certain study limitations and proposing avenues for future research. While our temporal analysis provides valuable information, enhancing the resolution through more frequent sampling intervals, especially during key regeneration phases, would offer a more nuanced understanding. Functional validation of identified genes and cellular dynamics is crucial, necessitating targeted experiments such as gene manipulations to confirm their roles in coral regeneration. Increasing sample size and incorporating biological replicates would enhance statistical robustness and reveal potential inter-individual variability. Integration of multi-omics data, including proteomics and metabolomics, holds promise for a comprehensive understanding of molecular processes. Considering the influence of environmental factors and exploring species-specific differences are vital for broader applicability. Investigating the elusive mechanisms governing cellular plasticity, signaling pathways during regeneration, and dynamic environmental influences will further deepen our comprehension. Overall, multi-omic integration, combined with a focus on species-specific variations, emerges as a promising approach for unraveling intricate regulatory networks and advancing coral regeneration research.

## Methods

### Sample collection and regeneration experiment

In the aquatic environment of the Xisha Islands (latitude: 15°40′−17°10′N, longitude: 111°−113°E), coral samples (*A. muricata*) were collected at a depth of approximately 5 to 10 meters. After collection, the corals were transferred to laboratory aquariums for a one-month acclimation period. Subsequently, three branches, each measuring 3−5 cm and exhibiting normal growth, were selected as biological replicates for the control group in the regeneration time-course experiment. Additionally, the tips of five branches (each not exceeding 1.5 cm) were excised for the preparation of cell suspensions in the control group for single-cell experiments, and one branch, measuring 3−5 cm and displaying normal growth, was chosen as the control group for the HRCT experiment.

Simultaneously, 47 branches measuring 3−5 cm had their apical tips (1 cm of the apical region) removed and set aside. Among these, 39 branches were designated as samples for the regeneration time series experiment. These were randomly selected at different time points after coral injury (3, 6, 9, 12, 15, 18, 21, 24, 27, 30, 33, 36, 39 days), with three samples taken at each time point to serve as biological replicates. Furthermore, five branches were selected for the single-cell experiment group, with their tips (not exceeding 1.5 cm) selected at 21 days post-injury. An additional three branches were reserved for HRCT 3D reconstruction experiments, with samples taken at days 14, 21, and 28 post-injury. Sampling was consistently performed around noon at 12:00.

The coral specimens were cultivated in a standard RedSea tank (redsea575, Red Sea Aquatics Ltd., London, UK) using the Berlin Method. The tank maintained a temperature of 26 °C, salinity at 1.025, and was equipped with specific devices, including three coral lamps (AI®, Red Sea Aquatics Ltd., London, UK), a protein skimmer (Reef Octopus Regal 250-S, Honya Co. Ltd., Shenzhen, China), a water chiller (tk1000, TECO Ltd., Port Louis, Mauritius), two wave devices (VorTech™ MP40, EcoTech Marine Ltd., Bethlehem, PA, USA) and a calcium reactor (Reef Octopus OCTO CalReact 200, Honya Co. Ltd., Shenzhen, China).

### HRCT and 3D reconstruction

We utilized a state-of-the-art 230-kV x-ray microfocus computed tomography system (Phoenix v|tome|x m, General Electric) at Yinghua NDT in Shanghai, China, to examine four *A. muricata* samples. Proprietary software from General Electric (GE) was employed to compile two-dimensional image reconstructions for each specimen based on matrices of scan slices. The derived slice data from the scans underwent analysis and manipulation using VG software. The construction of 3D reconstructions was performed using Mimics (v20.0) and VG Studio Max (v3.3.0), following a methodology described in a previous study(*60*).

### Single-cell suspension preparation, library construction, and sequencing

To prepare the single-cell suspension, a 0.035 M PBS solution with a pH of 8.0 was created by diluting 0.02 M PBS (Solarbio, P1010-2L, Beijing, China). The samples were thoroughly washed with this buffered solution, a process repeated three times. A digestion solution was then prepared by diluting 1‰ Tyrisin (Sigma-Aldrich, T1426-500MG, St. Louis, MO, USA) in the 0.035 M PBS solution. The obtained samples underwent a 2−3 hour digestion with this solution. Subsequently, the solution was filtered using 40 μm Fisherbrand^TM^ Sterile Cell Strainers (Thermo Fisher Scientific, 22-363-547, Waltham, MA, USA) and centrifuged at 1000 rpm for 5−6 minutes. The resulting supernatant was collected, and the digestion was terminated with 0.4 mg BSA (Sangon Biotech, A600332-0005, Shanghai, China) dissolving in 0.035 M PBS solution. Live cell counting and assessment of sample concentration and viability were performed using an automated cell counter (TC20^TM^, Bio-Rad Laboratories, Inc., Hercules, CA, USA). The qualified cells, after washing and resuspension, were prepared into a single-cell suspension at an appropriate concentration and then sent to Tianjin Novogene Bioinformatic Technology Co., Ltd. (Tianjin, China). In this company, library construction was carried out using the Chromium Controller (10x Genomics, Inc., Pleasanton, CA, USA), followed by quality control. Finally, sequencing was conducted on the Hiseq PE150 platform (Illumina, Inc., San Diego, CA, USA).

### mRNA extraction, library construction, and sequencing

For the regeneration time-course experiment, we designed 14 time points, including the normal state (0 days) and post-injury at days 3, 6, 9, 12, 15, 18, 21, 24, 27, 30, 33, 36, and 39. Each time point had three biological replicate samples, requiring a total of 42 Illumina cDNA libraries to be constructed (with mRNA > 1.5 µg for each sample). Additionally, at the normal state and the highly active repair period (day 21 post-injury), one PacBio cDNA library (with mRNA > 10 µg for each sample) and three miRNA libraries (with mRNA > 2 µg for each sample) were needed, respectively. For the construction of the PacBio cDNA library, an equal amount of mRNA from the corresponding three biological replicate samples was mixed together, totaling more than 10 µg.

During the mRNA extraction process, the mortar and pestle were consistently kept in liquid nitrogen to maintain sample integrity, strictly adhering to the manufacturer’s instructions. The mRNA extraction steps were as follows: 1) Grinding coral samples into granules; 2) Adding TRIzol® LS Reagent (Thermo Fisher Scientific, 10296028, Waltham, MA, USA) in a ratio of approximately 1:3 (sample: reagent) to the ground samples, continuing grinding to powders, and allowing natural freezing; 3) Continuing to add a small amount of TRIzol® LS Reagent and grinding until the sample completely dissolved, then transferring the liquid to a 50 mL centrifuge tube; 4) Centrifuging at 4°C and 3,000 rpm for 5−15 minutes, collecting the supernatant in a 50 mL centrifuge tube; 5) Adding BCP reagent (Molecular Research Center, BP 151, Cincinnati, OH, USA) in a ratio of 5:1 (sample: reagent) to the aforementioned centrifuge tube, shaking well, and letting it stand for 10 minutes; 6) Centrifuging at 4°C and 10,500 rpm for 15 minutes, collecting the supernatant, adding an equal volume of Isopropanol (Amresco, 0918-500ML, Radnor, PA, USA), mixing well, and incubating overnight at −20°C; 7) Centrifuging at 4°C and 10,500 rpm for 30 minutes, discarding the supernatant, and washing twice with diluted 75% Ice Ethyl alcohol, Pure (Sigma-Aldrich, E7023-500ML, Taufkirchen, München, Germany) to obtain the final mRNA product.

The samples were then sent to Tianjin Novogene Bioinformatic Technology Co., Ltd., for sequencing. PacBio Sequel II platform was used for full-length transcriptome sequencing, Illumina HiSeq X Ten platform was used for short-read transcriptome sequencing, and Illumina SE50 platform was used for miRNA sequencing. The company conducted quality control processing on the raw sequencing data.

### Reference genome alignment

The single-cell transcriptome sequencing data were aligned to the reference genome (BioProject: PRJNA544778) using Cell Ranger v6.0.0 software from 10x Genomics. Subsequently, gene expression quantification was performed to generate the cell-gene expression matrix. The clean reads from full-length transcriptome sequencing were mapped to the reference genome using GMAP v2017-06-20 software. For the processed small RNA reads with lengths ranging from 18 to 35 nucleotides, mapping to the reference genome and analysis were conducted using Bowtie v0.12.9. The reads mapped to the reference sequences were aligned to the miRBase database v22.0 using mirDeep2 v2.0.0.5 and srna-tools-cli to detect known mature miRNAs, and their expression levels and TPM values were quantified. Additionally, novel miRNAs were predicted through the integration of miREvo v1.1 and mirDeep2.

The clean reads from short-read transcriptome sequencing were aligned to the reference genome using HISAT v2.0.4. The Cufflinks v2.1.1 Reference Annotation Based Transcript (RABT) assembly method was employed to construct and identify both known and novel transcripts based on the HISAT alignment results. HTSeq v0.6.1 (-m union) was used for counting the read counts mapped to each gene. Subsequently, the mean transcripts per million (TPM) value for each gene across the three biological replicates was calculated, allowing for comparisons of gene expression levels between samples and normalization for differences in library size.

### Single-cell transcriptome analysis

The analysis and visualization of the cell-gene expression matrix were conducted using Seurat v4.9.9.9050 package in R v4.3.1 software. The distribution plot of all cell types within a specific cell type drew inspiration from the plot_attr function in the singleCellNet v0.1.0 package. The cell type distribution plots across different samples utilized the dittoBarPlot function from the dittoSeq v1.13.1 package. The FindMarkers function in the Seurat package was employed for differential expression analysis between two samples, considering genes with |avg_log2FC|>0.5 and adjusted p-value (p_val_adj)<0.1 as significantly DEGs. The clusterProfiler v4.8.3 package was used for GO enrichment analysis of significantly DEGs. Cell-cell communication analysis for the two samples was performed using the CellChat v1.6.1 package. Trajectory analysis for epidermal cells and calicoblasts was conducted using the monocle v2.28.0 package.

For functional annotation, gene sequences were aligned to the NCBI non-redundant (nr) database using Diamond v0.9.14.115 with parameters (diamond blastx--db nr--query input.fasta--out output.xml--outfmt 5--more-sensitive--max-target-seqs 20--evalue 1e-5). Protein functions were predicted using InterProScan v5.61-93.0 (./interproscan.sh-dp-iprlookup-goterms-pa-t p-i orfs.fasta-f xml) by analyzing protein structure domains and functional sites. Blast2GO v6.0.3 integrated the Diamond and InterProScan result files for final nr and GO annotation results. KEGG annotation was performed using KofamScan 1.3.0 (./exec_annotation-o kegg.out--cpu 8--format mapper-e 1e-5 orfs.fasta).

### Other sequencing data analysis

The clustering analysis of short-read sequencing data in the regeneration time-course experiment was performed using the TCseq v1.22.6 package and STEM v1.3.13 software. For full-length transcriptome, alternative splicing analysis utilized SUPPA v2.3 software, while alternative polyadenylation analysis employed TAPIS v1.2.1 software. Fusion gene identification criteria included alignment to two or more genomic loci in the reference genome (each aligned locus must be separated by at least 100kb), with a minimum of two supporting reads in the corresponding short-read data. Differential expression analysis of miRNA utilized the DESeq2 v1.38.3 package, considering genes with padj < 0.05 and |log2(fold change)| > 1 as significantly differentially expressed. Additionally, the prediction of miRNA target genes was carried out using the intersection of the miRanda v3.3a and RNAhybrid v2.0 software.

## Data availability

The scRNA-seq datasets generated in this study have been deposited in Figshare at https://doi.org/10.6084/m9.figshare.25107968 and the National Center for Biotechnology Information (NCBI) under BioSample accession numbers SAMN39649577-80. The time-course short-read RNA-Seq data, Pacbio full-length RNA-Seq data, and miRNA-seq data are available in the NCBI under BioSample accession numbers SAMN39649527-68, SAMN16675175-76, and SAMN39649571-76, respectively.

## Acknowledgements

We thank Dr.Jianming Zeng (University of Macau), and all the members of his bioinformatics team, biotrainee, for generously sharing their experience and codes. This work was supported by the Special Funding of Southeast University, grant number 6907038067, 6907038068, and 6907038148, and the Big Data Computing Center of Southeast University.

## Author contributions

T.H. and C. H. performed the experimental studies. T.H. carried out the computational analysis. T.H. participated in the writing of the paper. J.-Y. C. and Z.L. supervised the work. T.H., C. H., J.-Y. C., and Z.L. contributed to data discussion. All authors have read and approved the manuscript for publication.

## Supplementary Information

Supplementary_Figures.docx

Suplementary Dataset 1.xlsx

Suplementary Dataset 2.xlsx

Suplementary Dataset 3.xlsx

Suplementary Dataset 4.xlsx

Suplementary Dataset 5.xlsx

Suplementary Dataset 6.xlsx

Suplementary Dataset 7.xlsx

Suplementary Dataset 8.xlsx

Suplementary Dataset 9.xlsx

