## Supplementary material for "Unveiling the molecular choreography: In-depth single-cell transcriptomic exploration of the regenerative dynamics in stony coral": Suplementary Figures

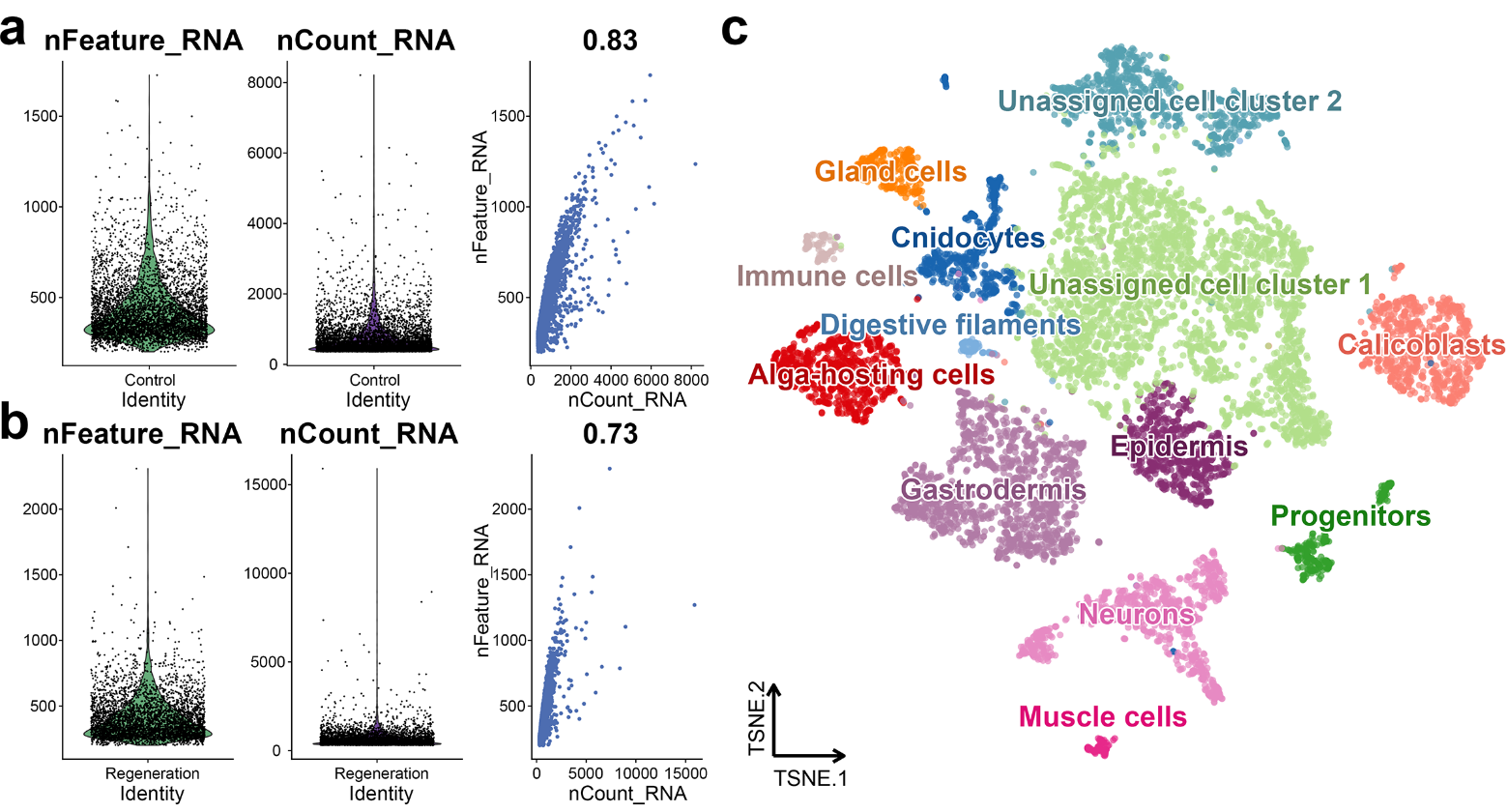

**Supplementary Fig. 1. Details of *A. muricata* cell atlas construction.** Visualization of QC metrics for healthy (referred to as 'Control' in **a**, and vigorously regenerating (referred to as 'Regeneration' in **b**, coral samples. **c**, The t-SNE map of *A. muricata*, where individual cells are depicted and distinguished by different colors based on their respective cell-type clusters.

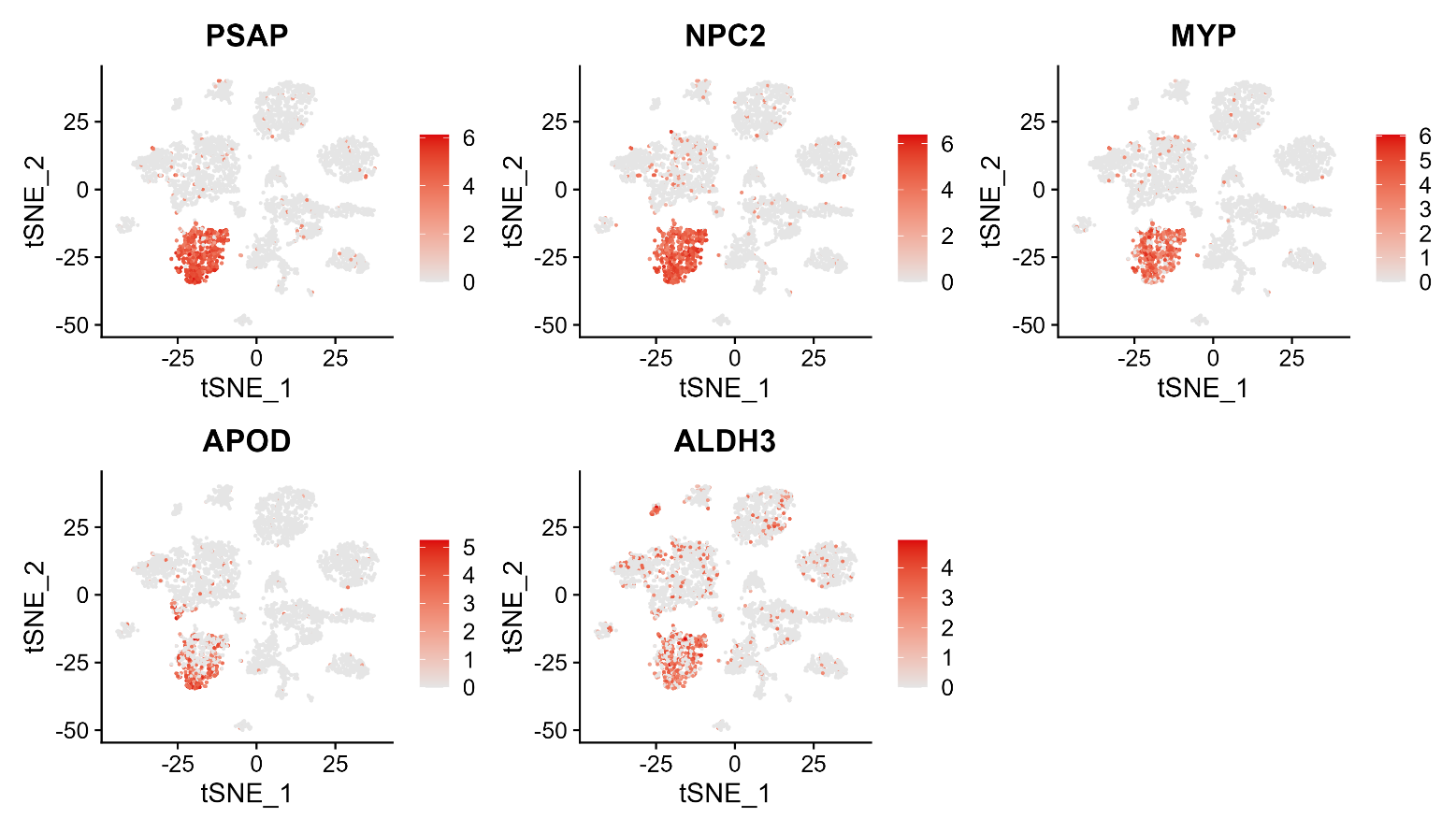
**Supplementary Fig. 2.** t-SNE visualization of the expression of marker genes in alga-hosting cells.

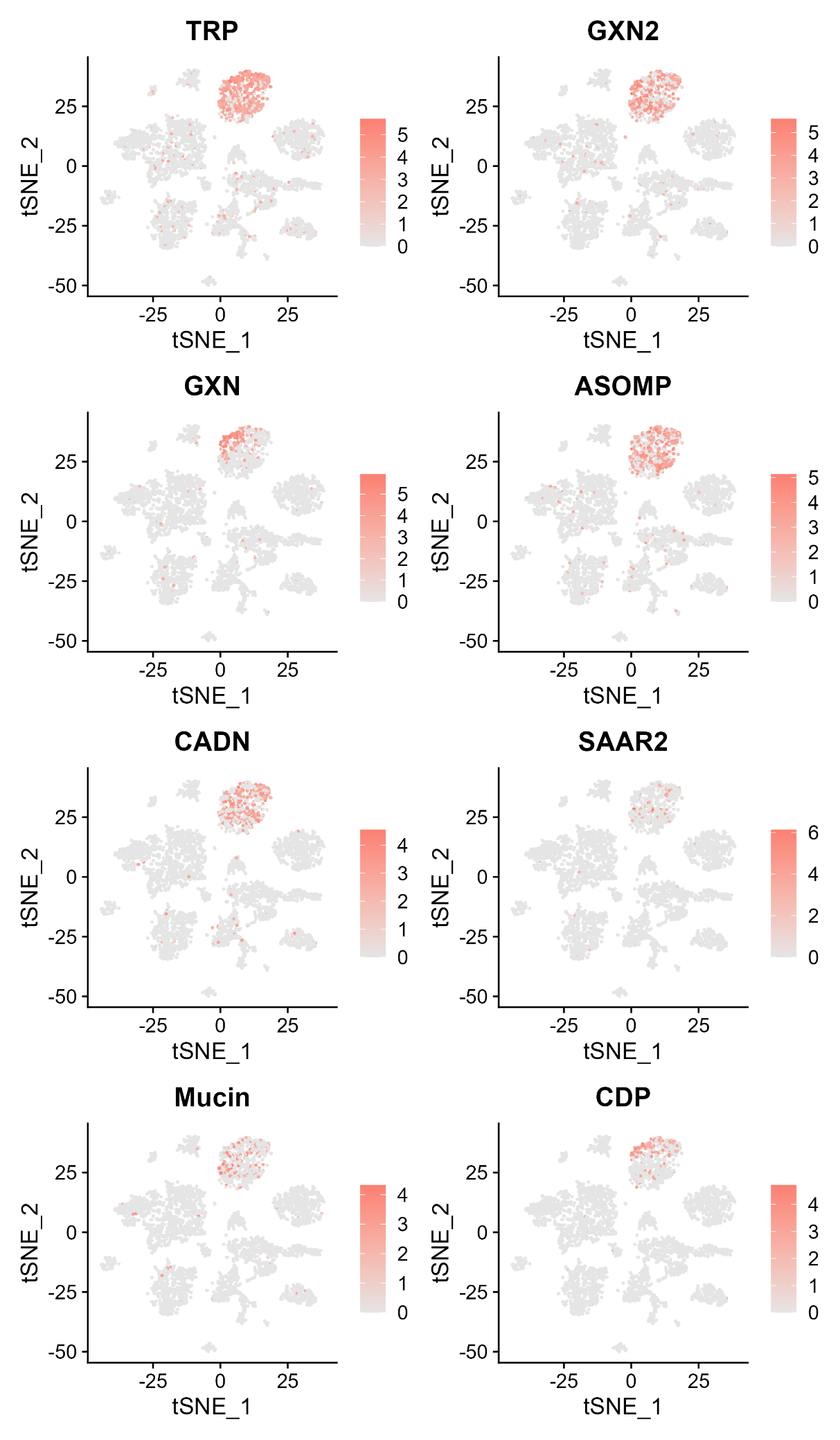
**Supplementary Fig. 3.** t-SNE visualization of the expression of marker genes in calicoblasts.

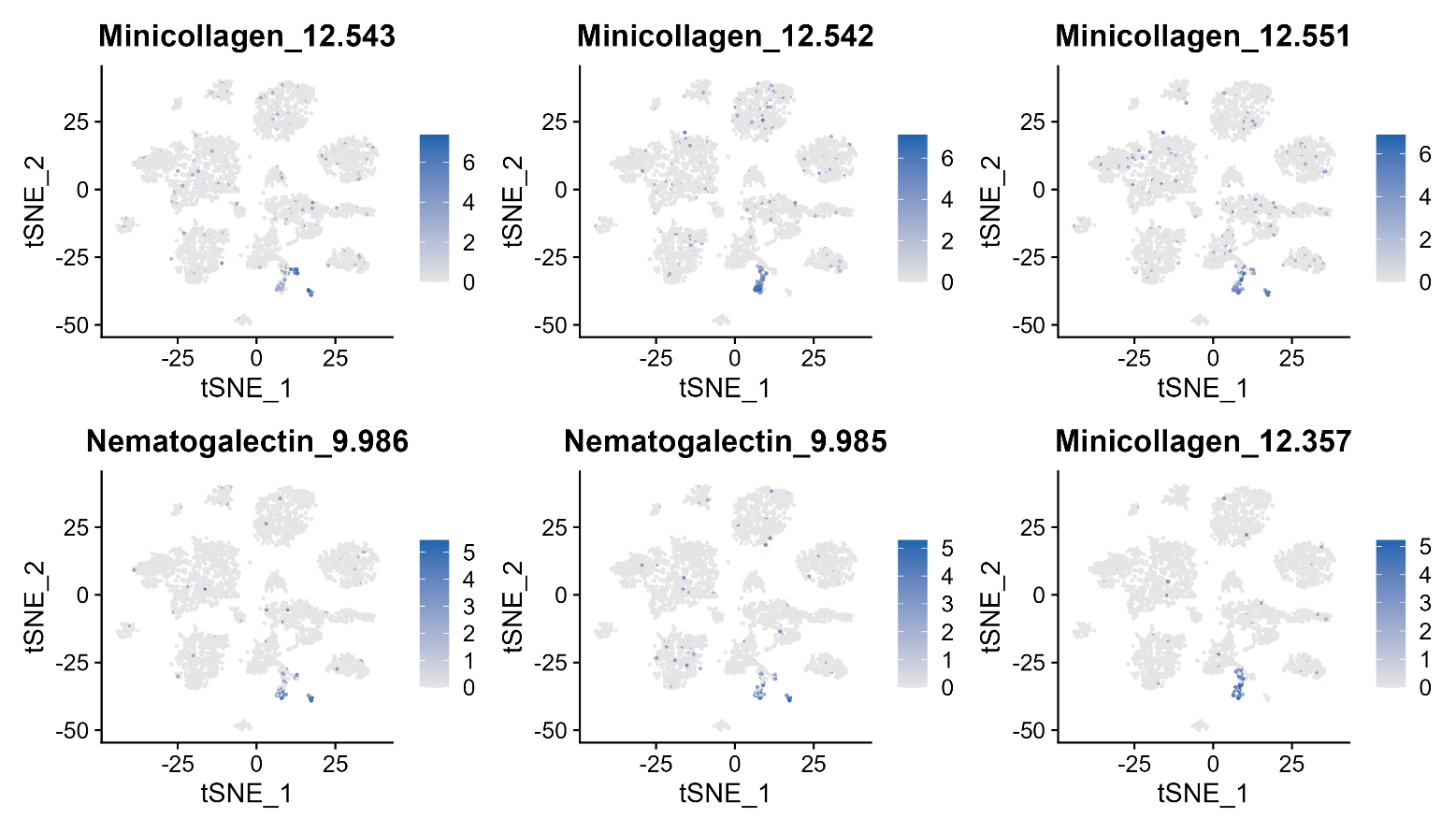
**Supplementary Fig. 4.** t-SNE visualization of the expression of marker genes in cnidocytes.

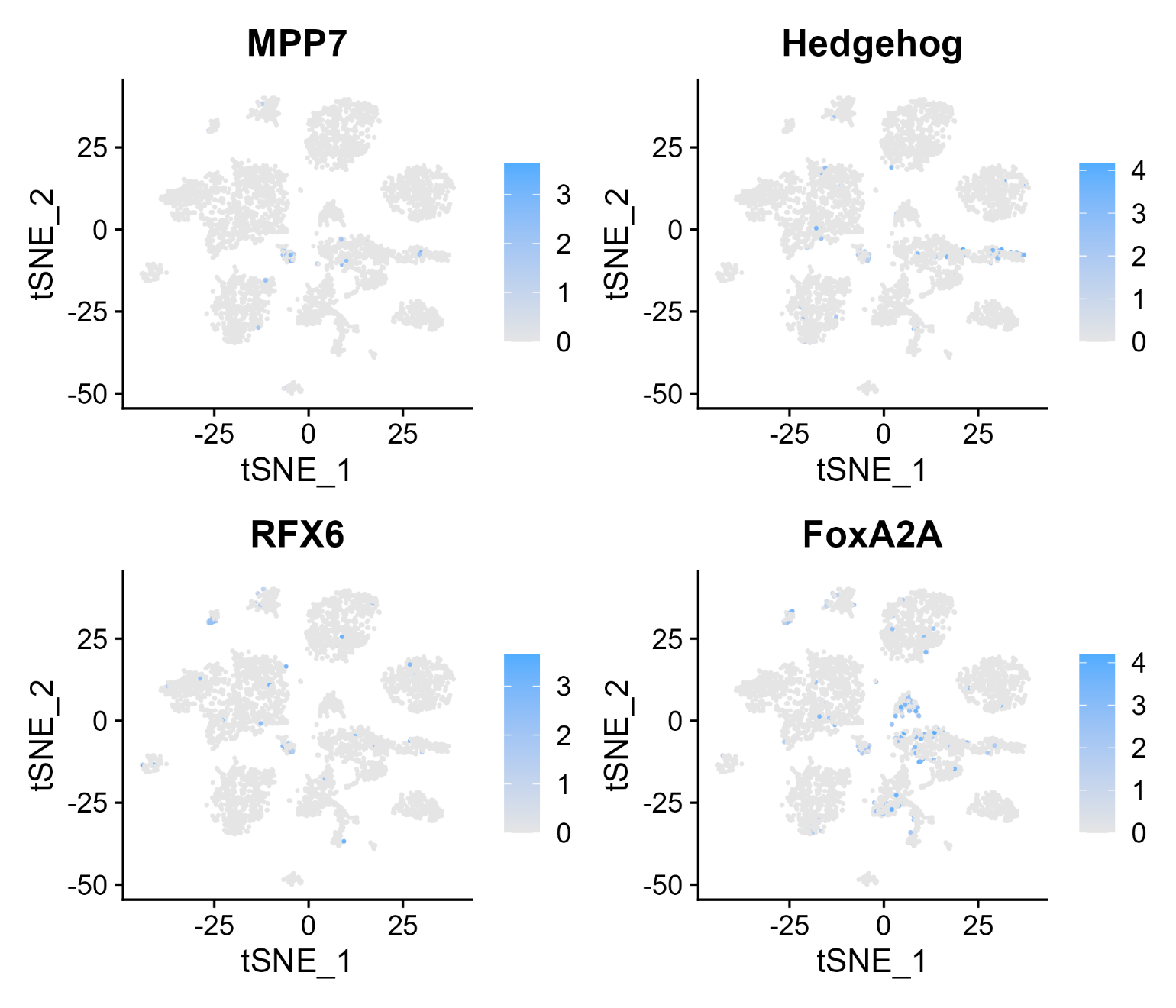
**Supplementary Fig. 5.** t-SNE visualization of the expression of marker genes in digestive filament cells.

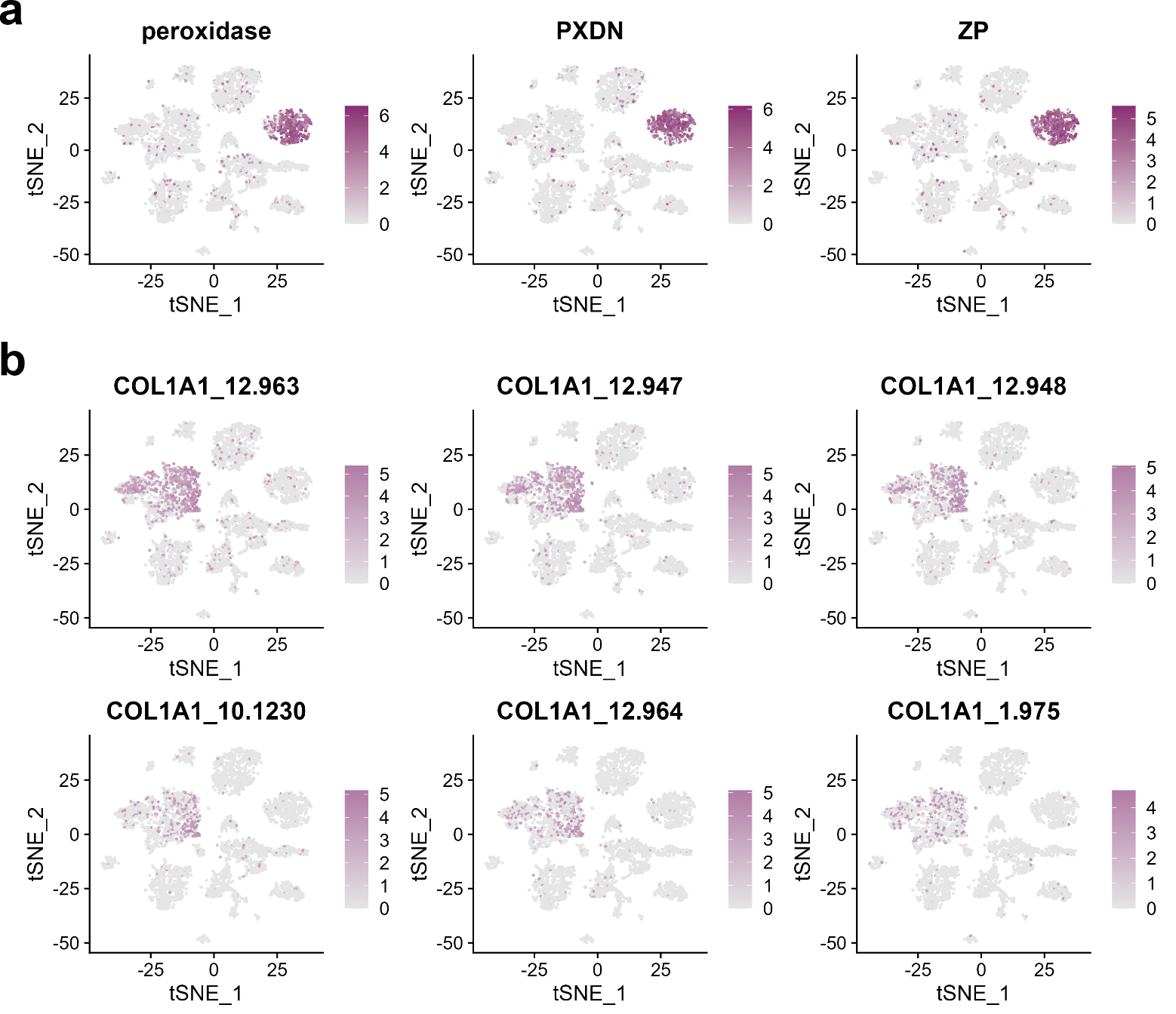
**Supplementary Fig. 6.** **a**, t-SNE visualization of the expression of marker genes in epidermal cells. **b**, t-SNE visualization of the expression of marker genes in gastrodermal cells.

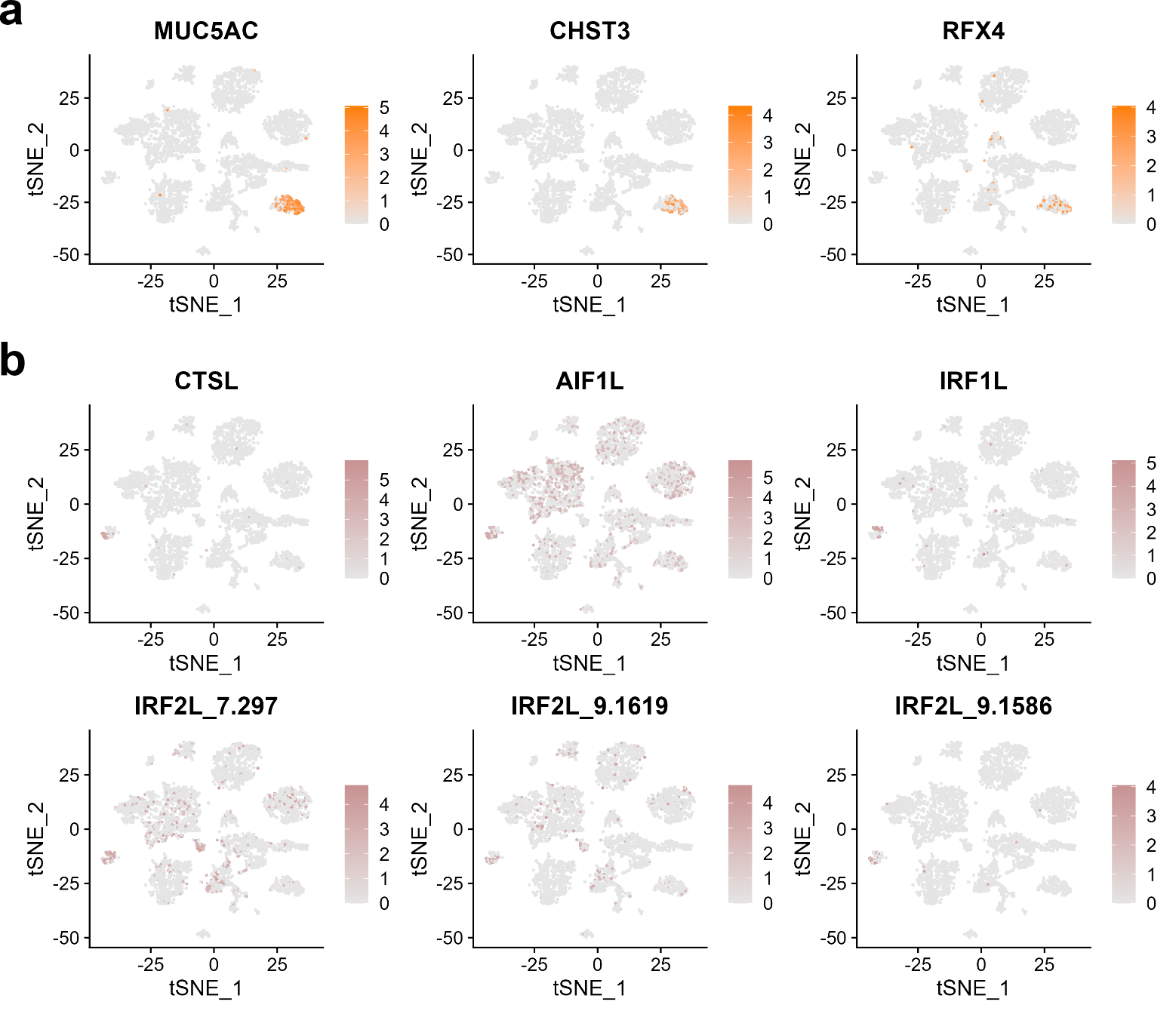
**Supplementary Fig. 7. a**, t-SNE visualization of the expression of marker genes in gland cells. **b**, t-SNE visualization of the expression of marker genes in immune cells.

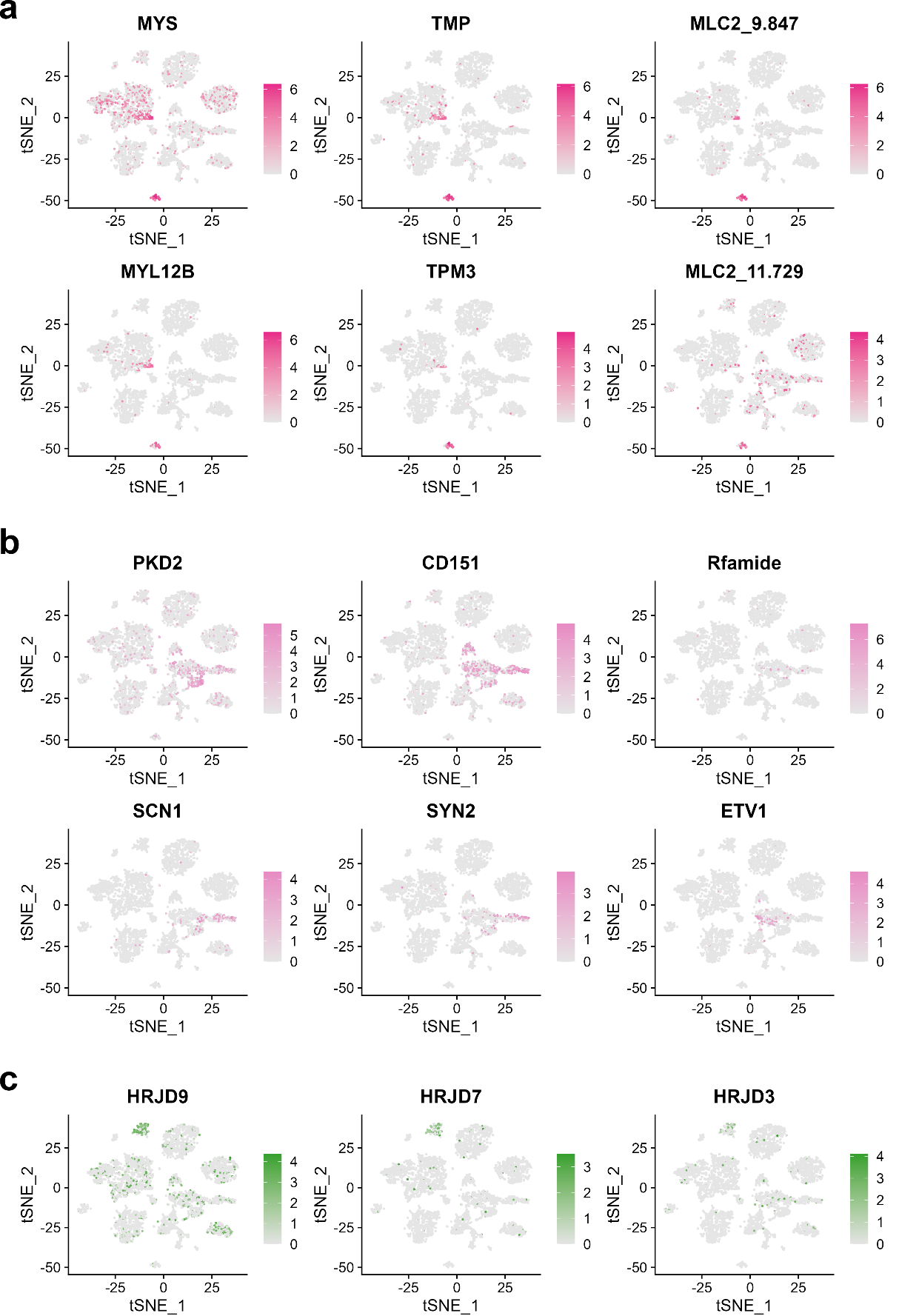
**Supplementary Fig. 8. a**, t-SNE visualization of the expression of marker genes in muscle cells. **b**, t-SNE visualization of the expression of marker genes in neurons. **c**, t-SNE visualization of the expression of marker genes in progenitor cells.

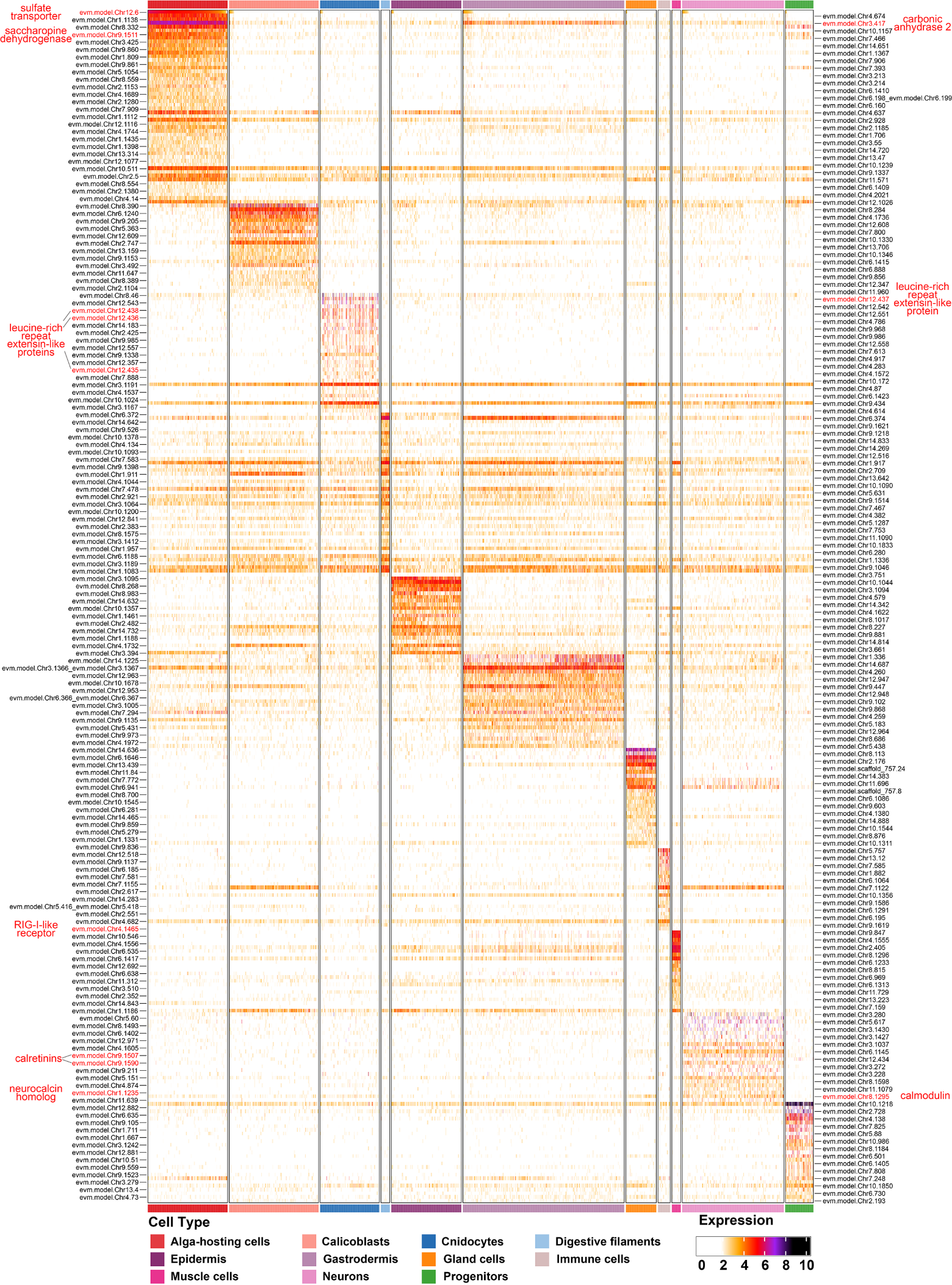
**Supplementary Fig. 9.** Candidate marker genes for different cell types.

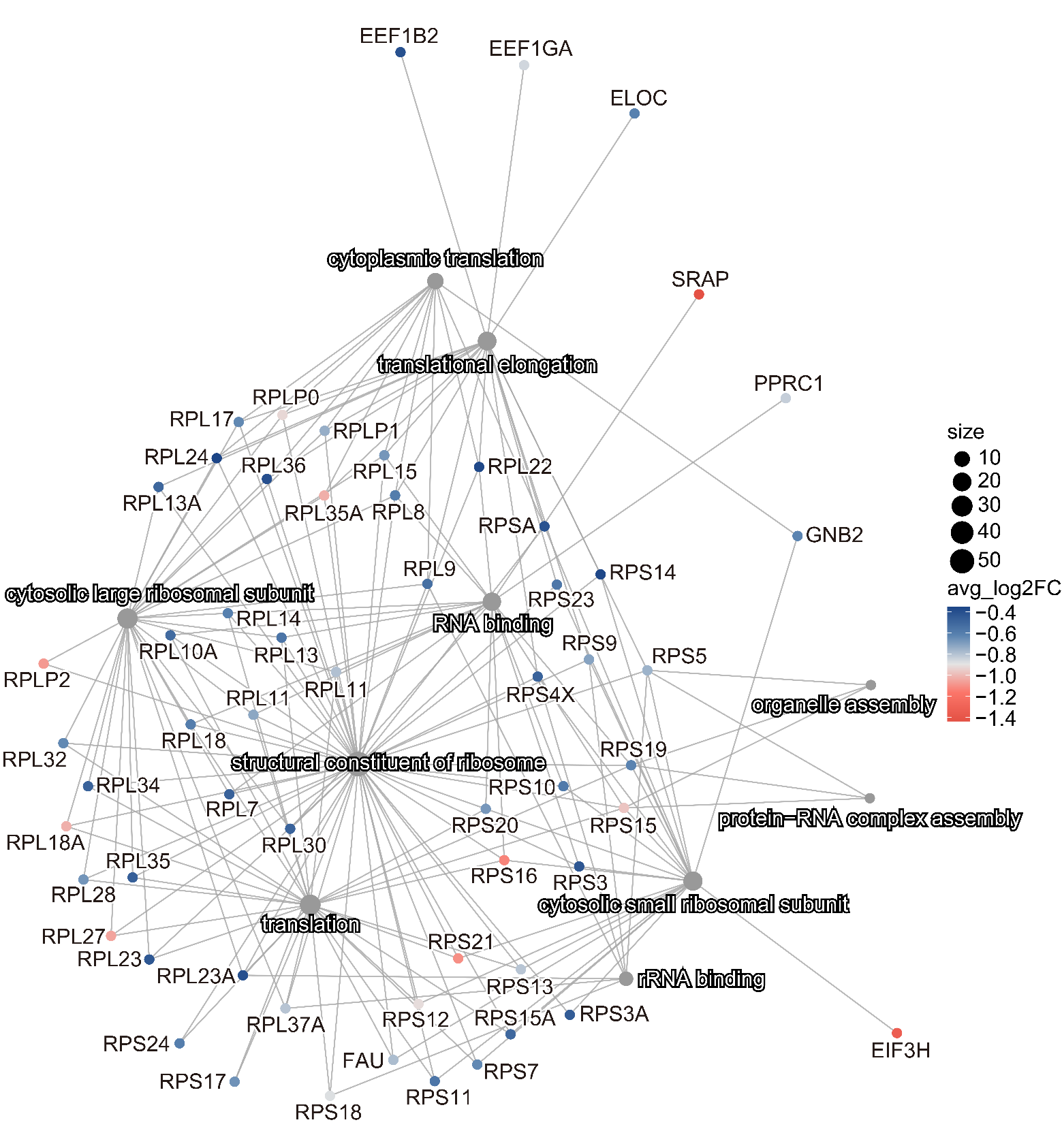
**Supplementary Fig. 10.** Cnetplot displays the remaining part of network linking significantly downregulated DEGs to enriched GO pathways.

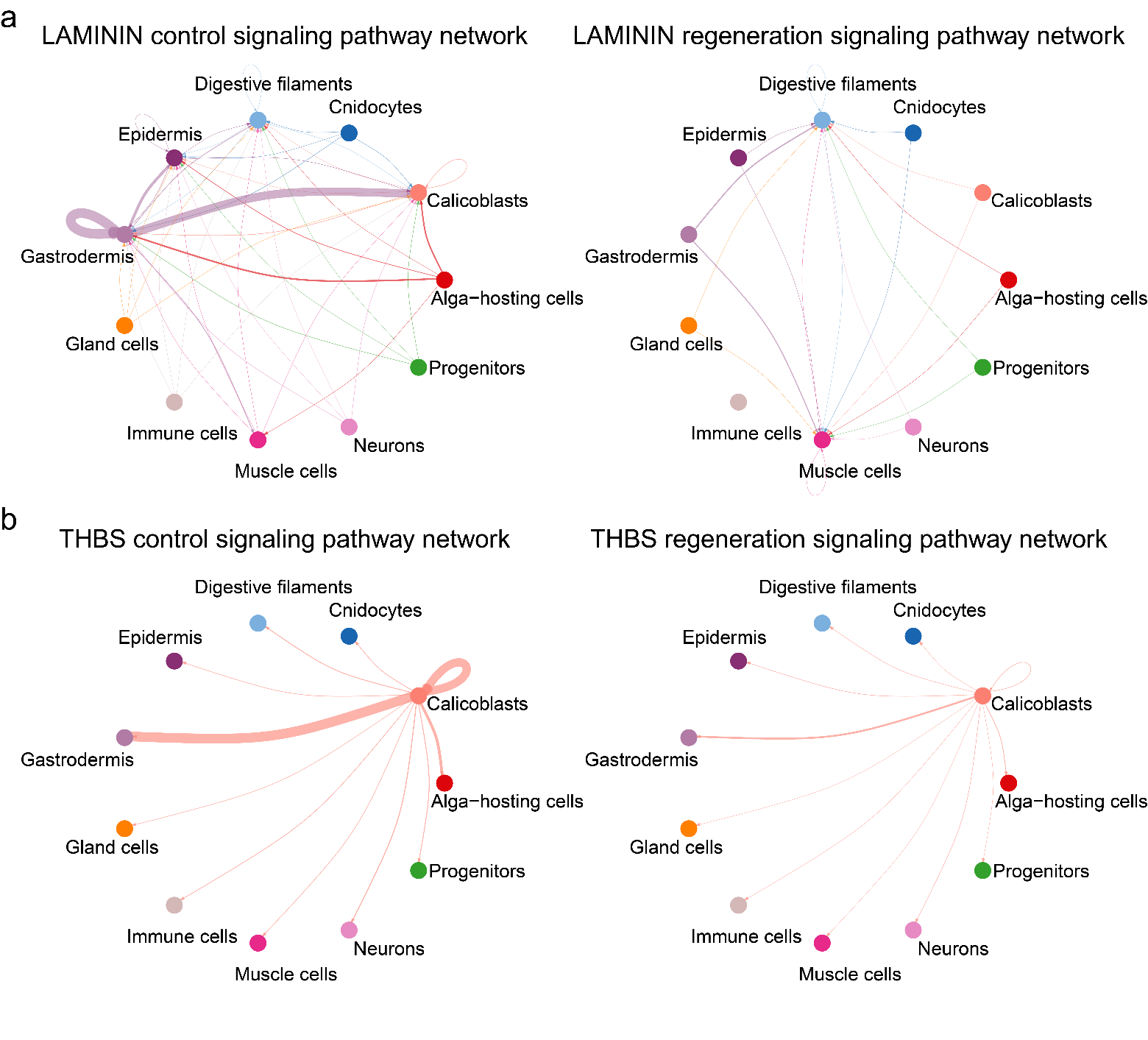
**Supplementary Fig. 11. a**, Pathway network of laminin signaling in different samples. **b**, Pathway network of THBS signaling in different samples. A thicker edge line indicates a stronger signal.

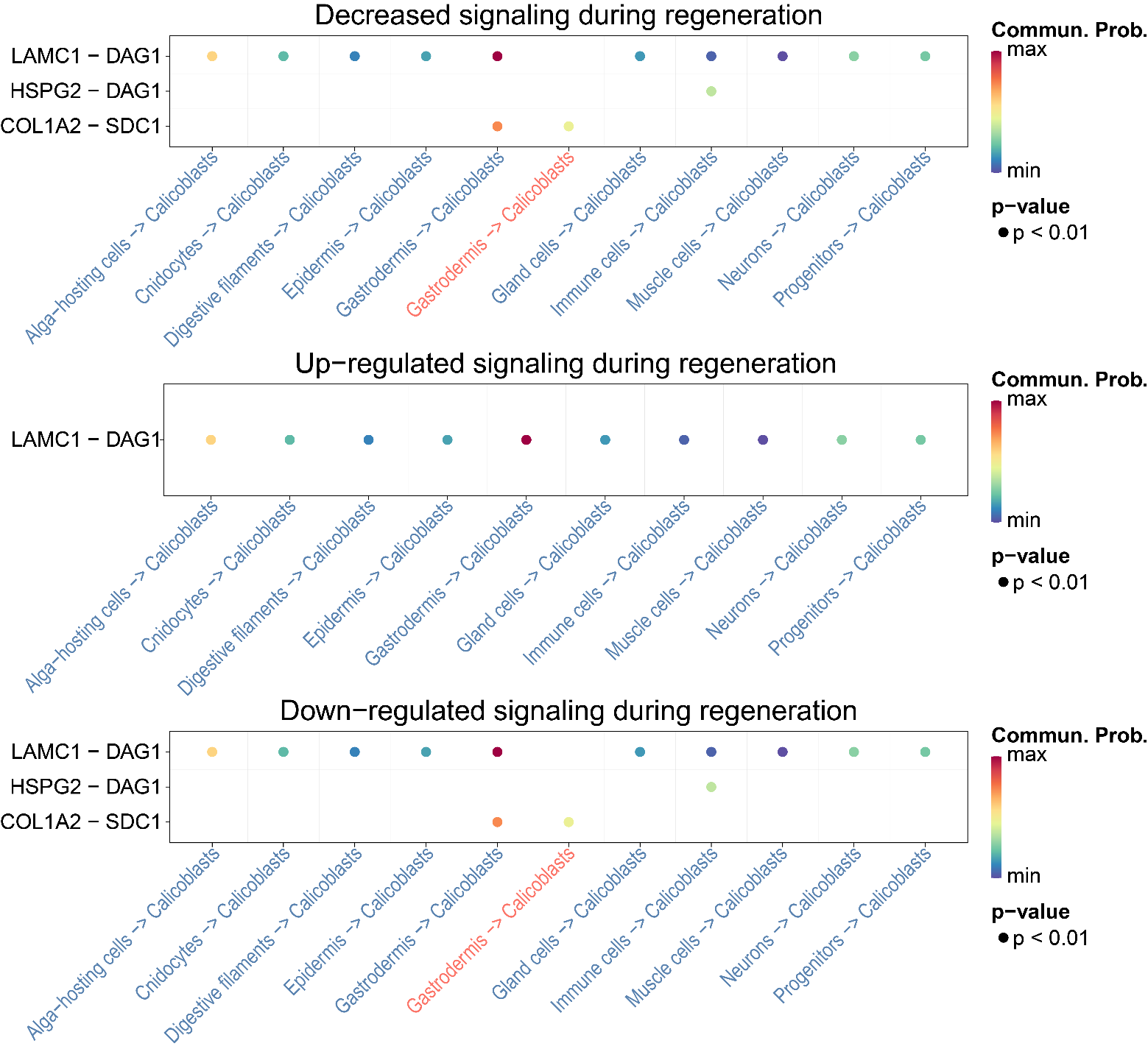
**Supplementary Fig. 12.** Increased (up-regulated) and decreased (down-regulated) in the communication probability of signaling ligand-receptor pairs from other cell types to calicoblasts in the regeneration sample compared to the control sample.

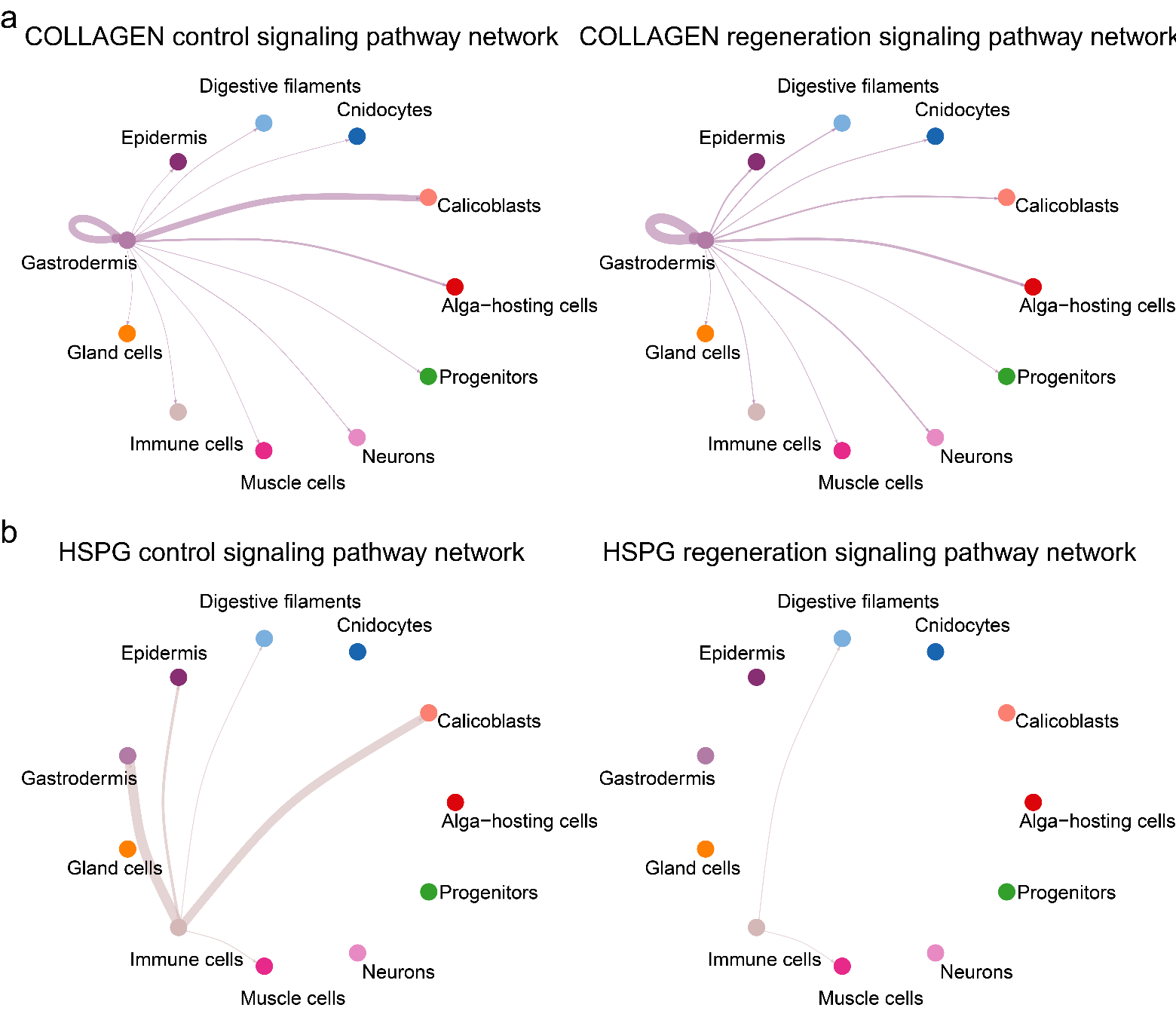
**Supplementary Fig. 13. a**, Pathway network of collagen signaling in different samples. **b**, Pathway network of HSPG signaling in different samples. A thicker edge line indicates a stronger signal.

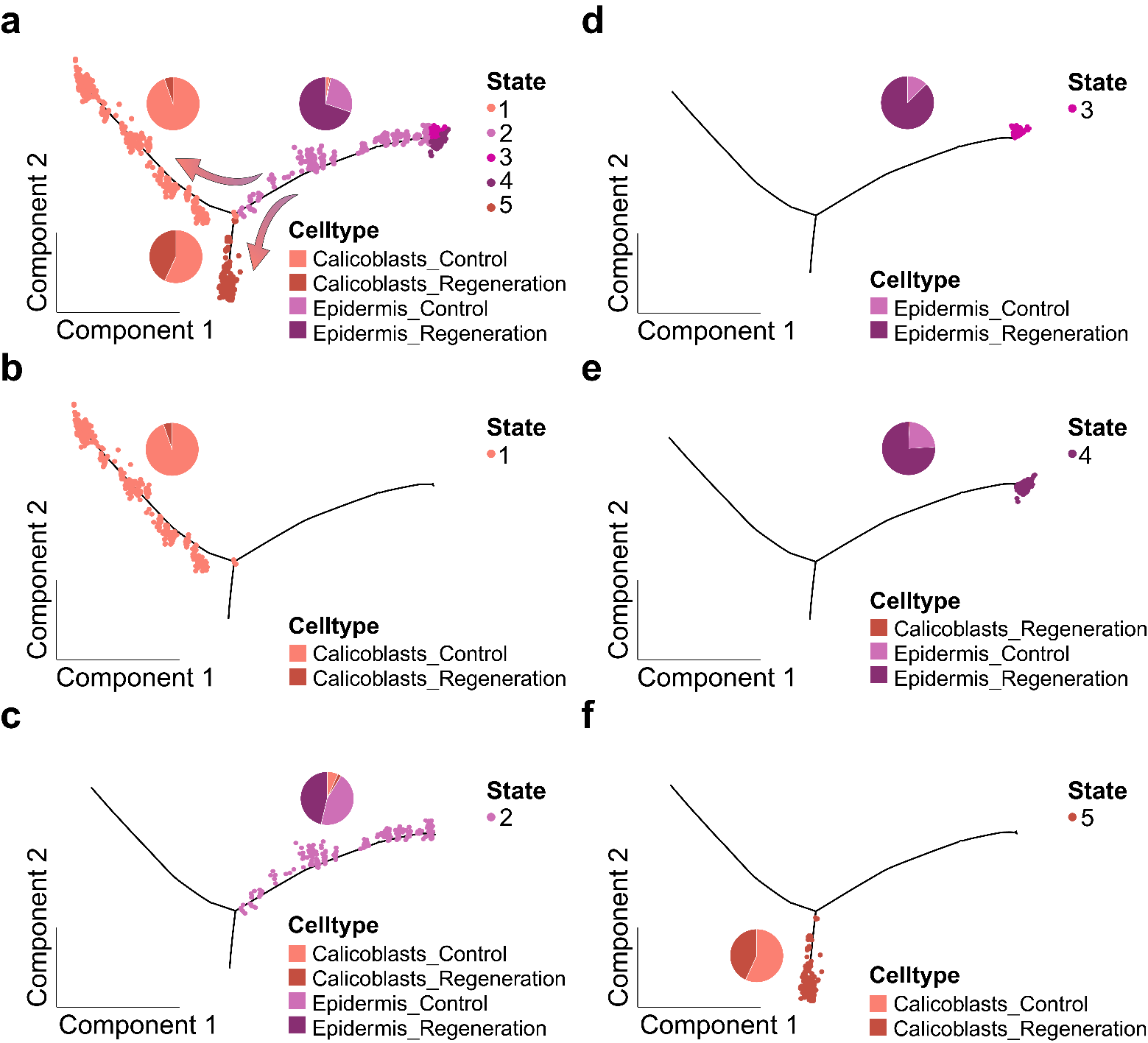
**Supplementary Fig. 14. Pseudotime trajectory showing different states.** Points along the trajectory are color-coded to represent cells belonging to different states. Pie charts depict the distribution of cells based on samples and cell types along different branches of the trajectory. **a**, All states. **b-f**, State 1-5.

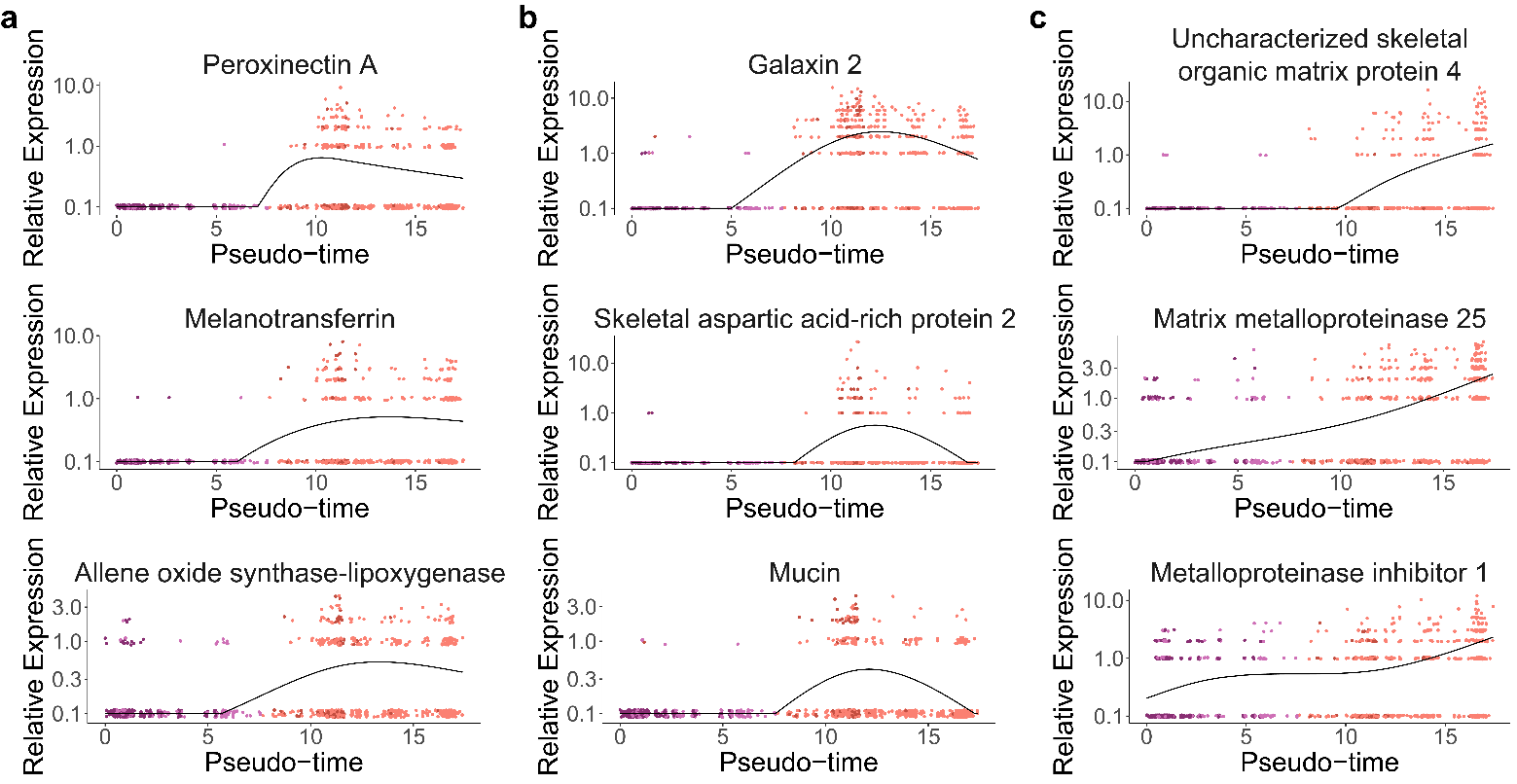
**Supplementary Fig. 15. Pseudotime analysis of other important genes based on cell types in Fig. 5e.** **a**, Genes associated with immune response in Cluster 1. **b**, Genes related to biomineralization in Cluster 3. **c**, Genes associated with biomineralization in Cluster 4.

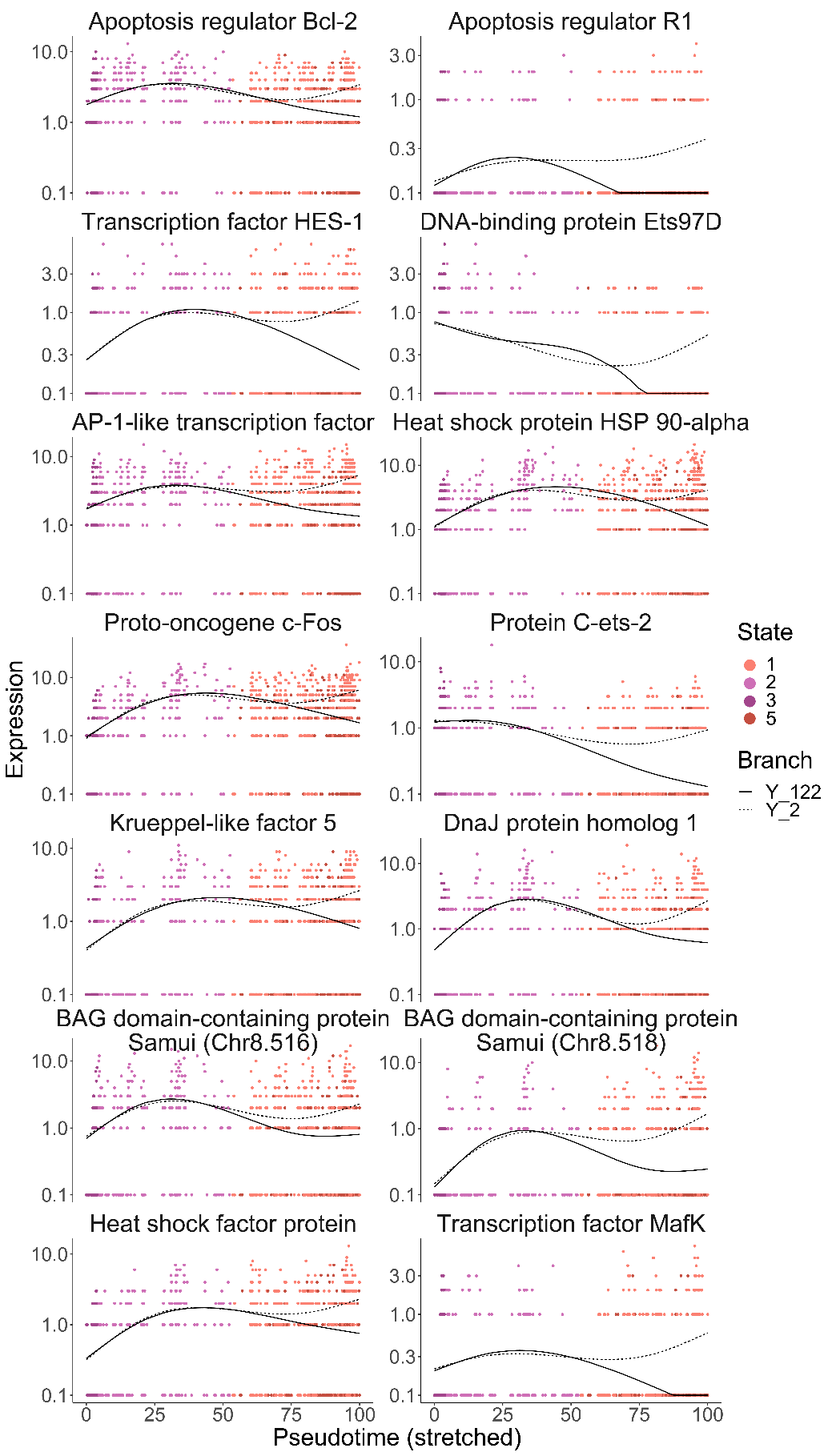
**Supplementary Fig. 16.** Differential expression analysis of genes enriched in two GO terms, protein stabilization, and apoptosis signaling pathways in Cluster 3 (labeled as "g" in Fig. 5), across different cell fates.

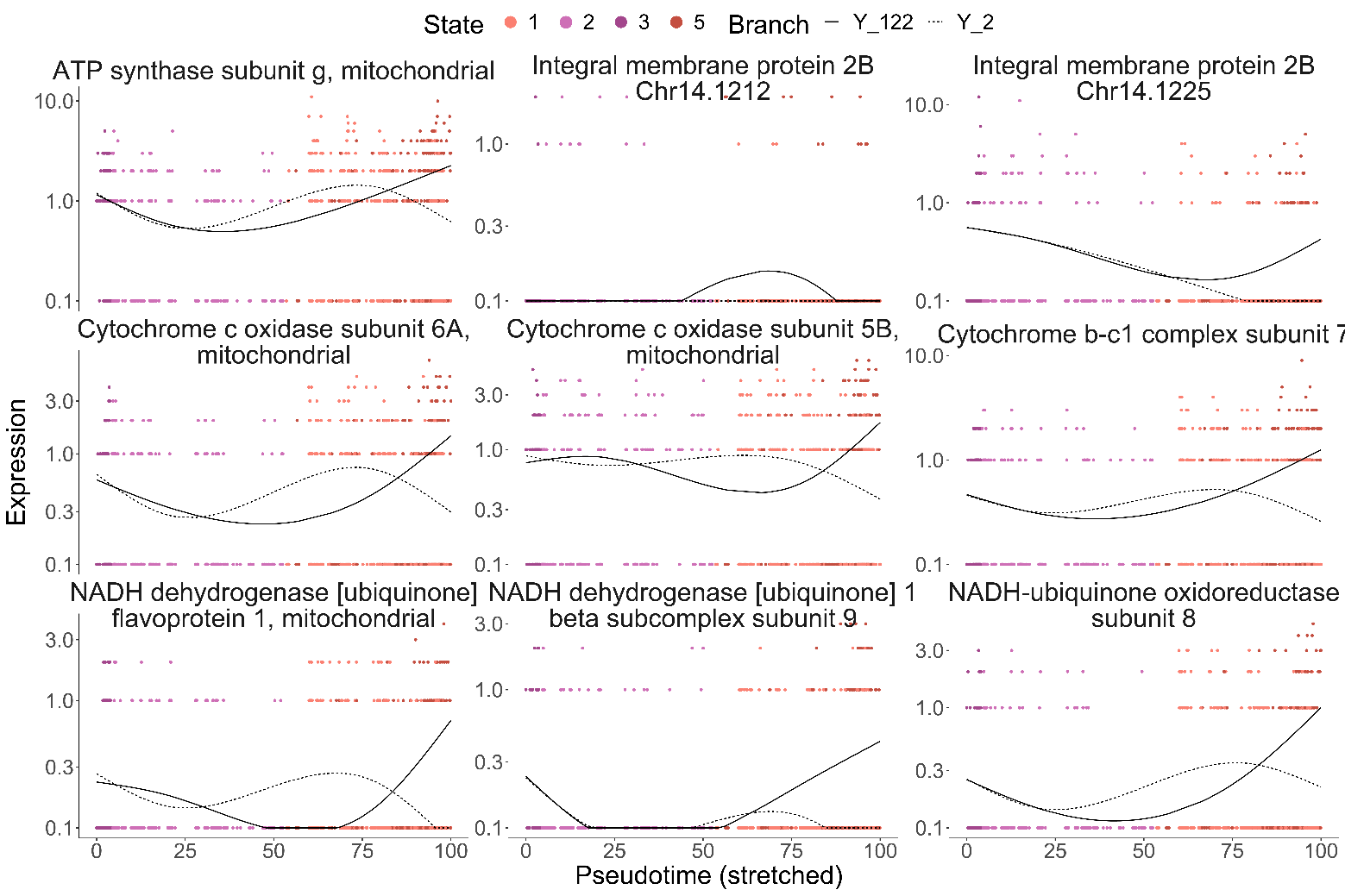
**Supplementary Fig. 17.** Differential expression analysis of genes enriched in three GO terms—proton motive force-driven ATP synthesis, transmembrane transporter complex, and regulation of cell population proliferation—in Cell fate 2 (labeled as "g" in Fig. 5), across various cell fates.

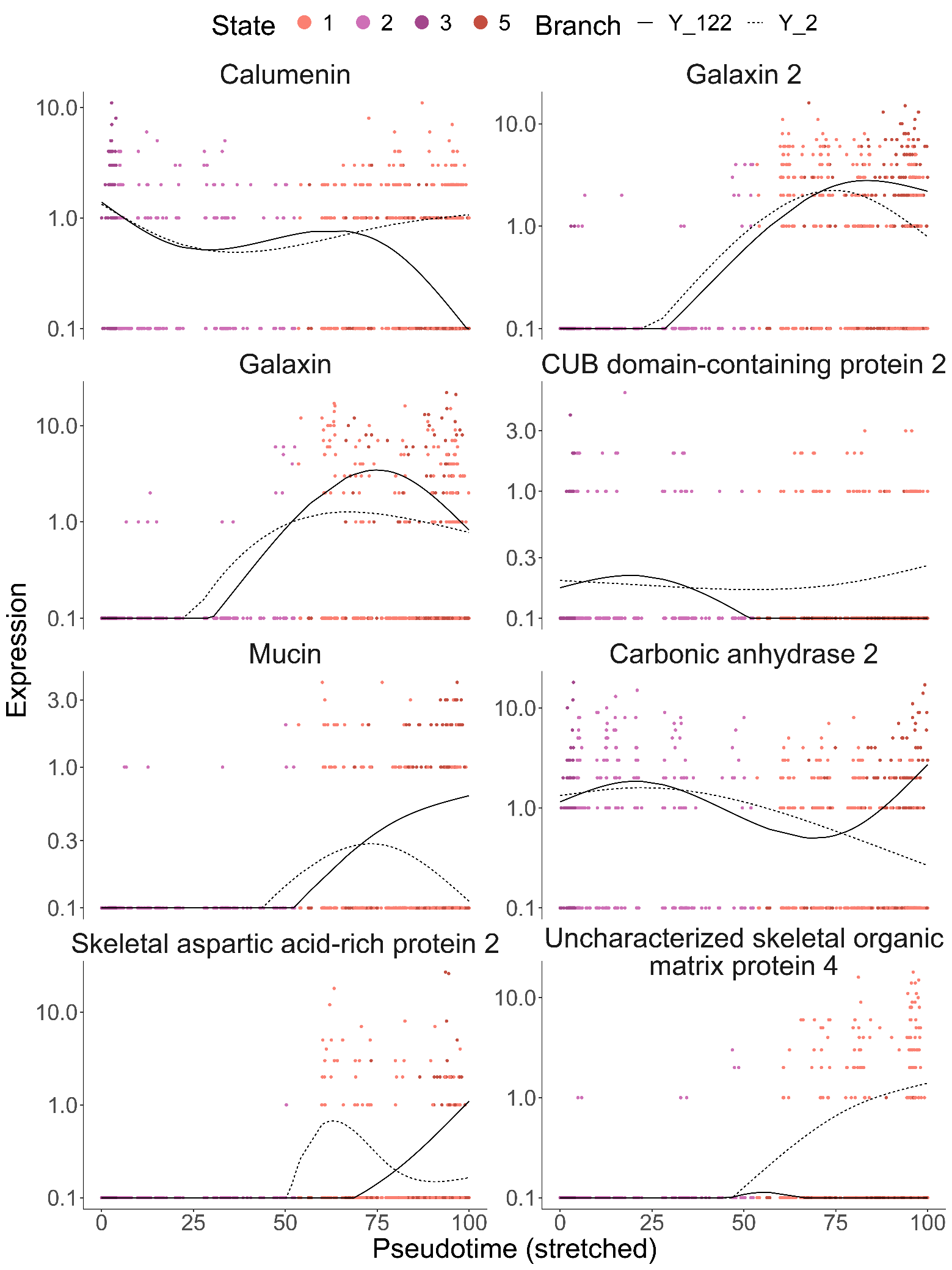
**Supplementary Fig. 18.** Differential expression analysis of genes related to biomineralization in different cell fates, as indicated in Fig. 5g.

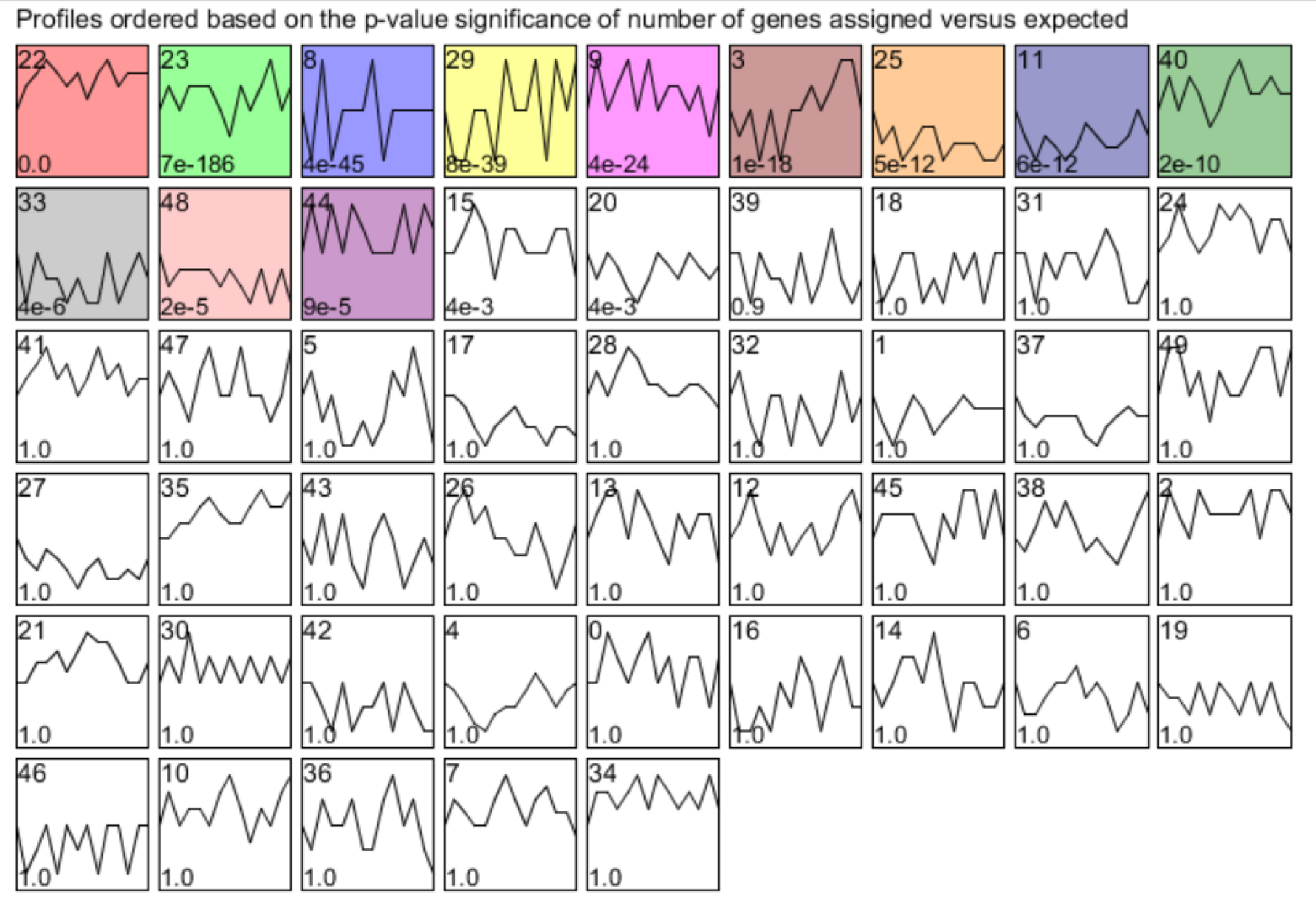
**Supplementary Fig. 19.** Clustering results obtained from temporal expression patterns of DEGs in bulk RNA-seq data by STEM.
